# Macroecological patterns in experimental microbial communities

**DOI:** 10.1101/2023.07.24.550281

**Authors:** William R. Shoemaker, Álvaro Sánchez, Jacopo Grilli

## Abstract

Ecology has historically benefited from the characterization of statistical patterns of biodiversity within and across communities, an approach known as macroecology. Within microbial ecology, macroecological approaches have identified universal patterns of diversity and abundance that can be captured by effective models. Experimentation has simultaneously played a crucial role, as the advent of high-replication community time-series has allowed researchers to investigate underlying ecological forces. However, there remains a gap between experiments performed in the laboratory and macroecological patterns documented in natural systems, as we do not know whether these patterns can be recapitulated in the lab and whether experimental manipulations produce macroecological effects. This work aims at bridging the gap between experimental ecology and macroecology. Using high-replication time-series, we demonstrate that microbial macroecological patterns observed in nature exist in a laboratory setting, despite controlled conditions, and can be unified under the Stochastic Logistic Model of growth (SLM). We find that demographic manipulations (e.g., migration) impact observed macroecological patterns. By modifying the SLM to incorporate said manipulations alongside experimental details (e.g., sampling), we obtain predictions that are consistent with macroecological outcomes. By combining high-replication experiments with ecological models, microbial macroecology can be viewed as a predictive discipline.

## Introduction

Microbial communities inhabit virtually every environment on Earth. Through their ubiquity, abundance, and diversity, microorganisms regulate the biogeochemical processes that sustain life on the biosphere. Furthermore, host-associated microbial communities play a crucial role in maintaining the health of many macroscopic forms of life, including humans, whose health is impacted by their microbiome [2, 3]. Given their environmental, medical, and economic importance, it is necessary to develop quantitative theories of ecology that allow researchers to explain, maintain, and alter the properties of microbial communities. A challenge of this scale is daunting, and the complexity of microbial communities has promoted the engagement of researchers from various disciplines with distinct approaches. Of these approaches, there are two that have substantially contributed towards our quantitative understanding of microbial communities in recent years: macroecology and experimental ecology.

Historically, the field of ecology has achieved considerable success by characterizing generalized patterns of biodiversity, an approach known as macroecology [4–10]. The macroecological approach is, fundamentally, statistical in nature, allowing for quantitative predictions and extrapolations to be made about the typical features of ecological communities without having to specify microscopic ecological forces. The generalized nature of this approach has allowed researchers to successfully characterize a diverse array of microbial ecological patterns [11–20] and spurred the development of mathematical models of microbial ecology grounded in statistical physics [21–29].

Through the macroecological approach, disparate patterns were recently unified by the observation that the typical microbial community follows three macroecological patterns: 1) the abundance of a given community member across communities follows a gamma distribution, 2) the mean abundance of a given community member is not independent of its variance (i.e., Taylor’s Law [30–32]), and 3) the mean abundance of a community member across communities follows a lognormal distribution [33]. These three general patterns can be captured by an intuitive mathematical model of density-dependent growth with environmental noise, the stochastic logistic model (SLM) of growth [33–35]. Building on its utility, the SLM has been successfully extended to quantitatively capture additional empirical microbial macroecological patterns. Examples that explicitly use the SLM include attempts to capture measures of ecological distances and dissimilarities between communities [36], alternative stable-states [37], correlations in abundance between community members [38], community-level measures across taxonomic and phylogenetic scales [39], and dynamics within and across human hosts at the sub-species level (i.e., strains) [19, 40]. The results of these studies demonstrate that a minimal mathematical model of ecological dynamics can capture a broad assemblage of microbial macroecological patterns.

The generality of macroecology is a useful feature that permits the construction of a working theory of microbial ecology [41, 42]. However, there is a trade-off between this generality and the ability of macroecology to provide causal connections between ecological mechanisms and observed patterns (e.g., the exponent of Taylor’s Law) [43]. This lack of causality is due to the inherent difficulty of manipulating macroecological patterns in natural communities. This study addresses this issue by expanding the scope of the macroecological approach from natural systems to experimental communities, providing the means to investigate how mechanistic manipulations alter macroecological patterns.

Experimentation has played a crucial role in the documentation and manipulation of ecological forces [44, 45]. The advent of 16S rRNA amplicon sequencing allows researchers to investigate the ecological dynamics of a large number of replicate microbial communities in a laboratory setting. This controlled approach has proven to be highly successful, allowing researchers to determine the extent that the assembly of microbial communities is reproducible and predictable [46–53]. Surprisingly, seemingly simple environments can maintain highly dissimilar microbial communities, even in systems harboring a single exchangeable resource [1]. This observation has led researchers to investigate the susceptibility of microbial communities to experimentally imposed ecological forces. A prominent example is the migration of individuals between communities, an ecological force that is amenable to experimental manipulation [54, 55] and can alter the heterogeneity of communities across replicates [1, 56, 57].

In this study we sought to bridge the gap between microbial macroecological patterns observed in nature and experiments performed in the lab. We consider experimentally assembled microbial communities exposed to qualitatively different forms of migration [1]. We first demonstrate that macroecological patterns can be recapitulated in a laboratory setting. By connecting these patterns to the predictions of the SLM, we developed a model of microbial macroecology that incorporates experimental details. Specifically, we focus on two migration treatments. The first treatment (referred to as regional migration) corresponds to a classical mainland-island scenario [58], where migrants from the community from which replicate communities were originally assembled (known as the progenitor community) continued to migrate over time. The second case, referred here as global migration, corresponds to a fully-connected metacommunity model [59], where migration occurs between communities that were assembled from the same progenitor community. We examine the ways in which the ecological force of migration can be manipulated, their resulting macroecological outcomes, and the degree that these outcomes can be predicted by the SLM. Using these results, we identify when and how the SLM succeeded in predicting the effects of experimental manipulation. By leveraging high-throughput ecological experiments alongside robust statistical patterns, we can strengthen the predictive and quantitative elements of microbial ecological theory.

## Materials and methods

### Experimental data

We identified an appropriate dataset to investigate the macroecology of experimental microbial communities. A recent experimental study maintained ∼ 100 replicate communities assembled from a single progenitor soil community and imposed a variety of demographic treatments [1]. Here, a given microcosm was inoculated from a single progenitor community. All microcosms contained a single supplied carbon source (i.e., glucose). Each community was allowed to grow for 48 hours. A small fraction of the volume (i.e., an aliquot) was then sampled and used to inoculate a new microcosm with the same profile of supplied resources. This procedure of inoculating a microcosm with an aliquot and allowing it to grow for a given period of time is known as a transfer cycle (Fig. 1a). Transfer cycles were repeated 18 times per replicate community.

**Figure 1.**
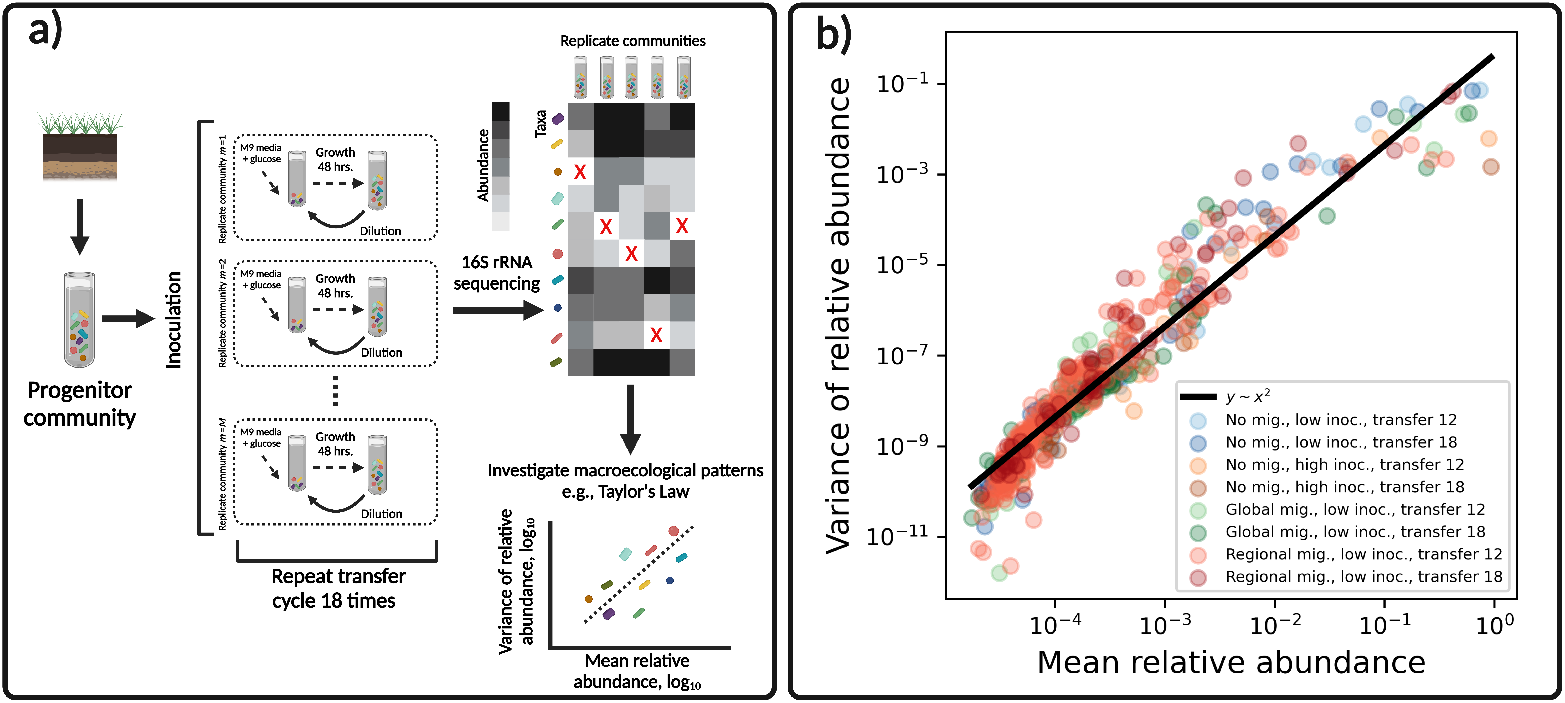
Emergent variation in experimental communities. **a)** Microbial community assembly experiments provide a means to maintain diversity in a laboratory setting. These experiments are commonly performed by growing a community sampled from a given environment (e.g., soil) and using it to inoculate a large number of replicate microcosms containing the same resource (e.g., M9 minimal media with glucose). These replicate communities are allowed to grow for a period of time (e.g., 48 h.) before aliquots are taken and diluted into microcosms containing replenished resource. This process known as a “transfer cycle” is repeated a given number of times (e.g., 18) and the resulting communities are sequenced via 16S rRNA sequencing. The abundances of community members have the potential to provide the variation necessary to investigate the existence of macroecological patterns in an experimental setting (e.g., Taylor’s Law), a goal of this study. Red “X” symbols represent the absence of a community member in a sample. **b)** Variation in abundance consistently arises across treatments and timescales in experimental communities. This variation can be captured by the relationship between the mean and variance of relative abundance, a pattern known as Taylor’s Law. Each data point represents statistical moments calculated across replicate communities for a single ASV from a single experimental treatment.

The demography of replicate communities was systematically altered by manipulating the form of migration. For the first manipulation, aliquots of the progenitor community was mixed with aliquots from the previous transfer cycle, so that the community composition of a microcosm at the start of a new transfer cycle was comprised of both the composition of the progenitor and the previous transfer cycle. This manipulation is referred to as regional migration. In the second case, migration was manipulated by sampling aliquots of each community at the end of a given transfer cycle. These aliquots were mixed and redistributed at the start of the subsequent transfer cycle. This manipulation is referred to as global migration. Migration manipulations were only performed for the first 12 dilution cycles (out of the total 18 transfer cycles). This experimental design provides a series of time points where migration was and was not performed within each replicate community (Fig. 3a).

A number of communities were also inoculated with a large aliquot of the progenitor community. Macroecological patterns of these communities were examined but were excluded from subsequent analyses since this manipulation only occurred at the first transfer cycle and did not induce lasting macroecological effects. Community data for this experiment was obtained via 16S rRNA sequencing. We reprocessed all raw FASTQ data from the original study to obtain Amplicon Sequence Variants (ASVs) using the package DADA2 [60]. Table S1 reports the number of replicate communities sampled at each transfer. S1 Text summarizes technical details about the experiment, which can can also be found in the original manuscript [1].

### The macroecological predictions of the stochastic logistic model

Populations at low abundance often grow at an initially high rate that proceeds to decrease as they approach the maximum abundance that an environment can support. This is the central idea underlying logistic growth and it can be captured by two parameters: 1) the minimal length of time required to reproduce (inverse of the maximum growth rate), *τ* and 2) the carrying capacity of an environment, *K*. This simple picture assumes a fully deterministic environment. Environmental stochasticity can be added as fluctuations in the growth rate, where the strength of environmental noise is controlled by the parameter *σ*, representing the coefficient of variation of growth rate fluctuations. The Stochastic Logistic Model (SLM) captures this two fundamental ecological processes under the form of a Langevin equation [61, 62]

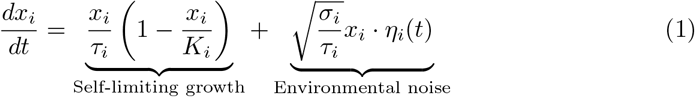

where *x*_*i*_ is the relative abundance of ASV *i*. The term *η*_*i*_(*t*) introduces stochasticity into the equation. It is defines as white noise: the expected value of *η*(*t*) is *η*(*t*) = 0 and the time correlation is a delta function ⟨*η*(*t*)*η*(*t*^*′*^) ⟩ = *δ*(*t − t*′) (implying that the noise at time *t* is uncorrelated with the noise at time *t*′). This model can be extended to time-correlated environmental fluctuations [33].

The SLM reproduces the three empirical macroecological patterns that are found in microbial communities across natural environments: 1) gamma distributed abundances, 2) a power law relationship between mean relative abundance and the variance of relative abundance (i.e., Taylor’s Law), and 3) a lognormally distributed distribution of mean relative abundances [33]. To provide the necessary context for the study, we briefly summarize these three patterns and their connection to the SLM.

#### The Abundance Fluctuation Distribution

The Langevin equation defined in Eq. 1) defines a stochastic trajectory *x*_*i*_(*t*). The probability *P* (*x*_*i*_,*t*|*x*_*i*_(0), 0) that the abundance of species *i* equals *x*_*i*_(*t*) at time *t* when starting from initial condition *x*(0) at time *t* = 0 can be obtained from the Fokker-Planck equation corresponding to Eq. 1 [61]. For large times, this probability converges to the stationary probability distribution *P* ^*∗*^(*x*_*i*_), which in the case of SLM takes the form of a Gamma distribution

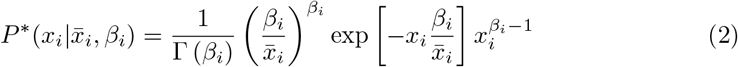

where the two parameters 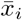 and *β*_*i*_ can be expressed in terms of the statistical moments of the distribution and also in terms of parameters appearing in Eq. 1

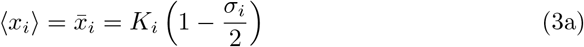

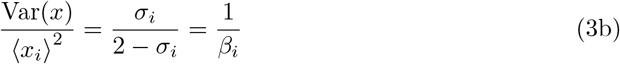

The expression above hold for absolute abundances, while we have access to finite number of reads obtained by sequencing, which are affected by both compositionality and sampling effects. We can model our sampling process as the Poisson limit of a multinomial distribution, obtaining a form of the AFD that explicitly accounts for the effect of sampling [33]. Using this distribution, we can calculate the probability of obtaining *n* reads out of a total sampling depth *N* for the *i*th ASV as

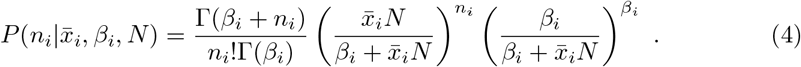

Often in ecology we are interested in the relationship between the mean relative abundance of a community member and the fraction of communities where it is present (i.e., its occupancy) [14, 63, 64]. We can derive a prediction of occupancy using the sampling form of the AFD by setting *n* = 0 and noticing that probability that an ASV is present is the complement of its absence. Averaging over *M* communities, one obtains a prediction of occupancy

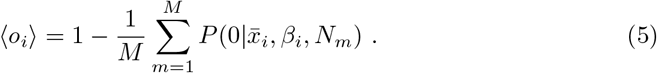

Prediction error was estimated as described in Text S2.

#### Taylor’s Law

Taylor’s Law describes the empirical relationship between the mean and variance of the relative abundance [30, 31] calculated across ASVs

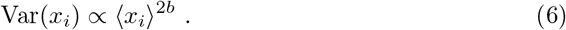

By comparing this expression to eq. 3 one can see that the case where 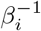 is independent of 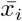 corresponds to *b* = 1. In the context of the SLM, this choice implies that the strength of environmental noise is constant for all ASVs (*σ*_*i*_ = *σ*). Estimates of *b* were obtained by log transforming both axes and performing ordinary least squares regression with `SciPy v1.4.1`.

#### The Lognormally distributed Mean Abundance Distribution

In observed data, the mean relative abundances (calculated either across communities or over time) of ASVs, known as the Mean Abundance Distribution (MAD), frequently follows a lognormal probability distribution

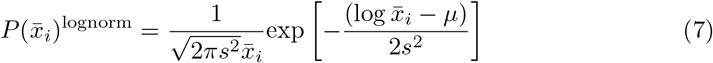

To account for sampling effects, we use a modified form of the lognormal that considers mean abundances with a coverage greater than

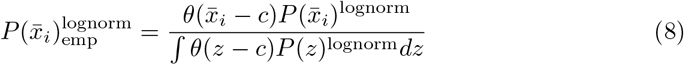

where *θ*(·) is the Heaviside step function. Parameters of this modified lognormal were fit to the empirical MAD as previously described (see Supplementary Note 7 in [33]).

### Incorporating migration into the SLM

The original form of the SLM is not directly applicable to microbial community assembly experiments. As discussed above, such experiments are performed as transfer cycles, where growth occurs over a period of time within a microcosm before an aliquot is transferred to an environment with replenished resources. Phenomenological models of logistic growth are appropriate for modeling dynamics within a single transfer cycle under certain conditions (S3 Text). In addition, the time-dependent form of Eq. 9 has been derived (i.e., *P* (*x*_*i*_|(*t*), *t x*_*i*_(0), 0) and could be used to model the temporal dynamics within a given transfer cycle [65] (description of solution in S5 Text). However, no attempt has been made to incorporate experimental details into the SLM so that it can be used to obtain predictions for the macroecological effects of experimental manipulations.

First, we extended the SLM to incorporate the transfer cycle process. We consider a piece-wise form of the SLM to describe the dynamics within the *k*th transfer cycle

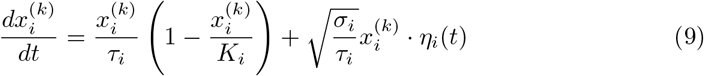

where the dynamics start at *t* = 0 and continue until time *t* = *T*. At time *T* an aliquot of the community is sampled. The volume of the aliquot relative to the total volume of the microcosm is known as the transfer dilution rate *D*_transfer_. If the total abundance of the community is known at time *T*, then the total abundance at the start of the next transfer cycle can be calculated as *N* ^(*k*)^(0) = *D*_transfer_ ·*N* ^(*k−*1)^(*T*). The process of sampling community members at the end of a transfer cycle can be modeled as multinomial sampling process (Text S6). With this sampling process and our piece-wise form of the SLM, we have a minimal dynamical model of a microbial community assembly experiment.

While they represent different forms of migration, both treatments are similar in that they are implemented by altering the initial abundance of a community member at the start of a transfer cycle. We can formulate the number of cells sampled due to migration at the start of a transfer cycle for a given ASV as follows

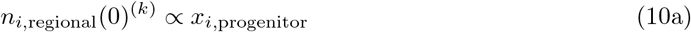

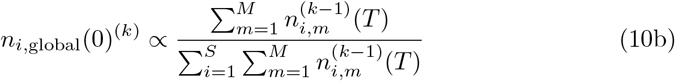

The sampling process of migrants can again be modeled using a multinomial distribution (S6 Text). We can now examine how the mean initial *relative* abundance of a given ASV depends on migration. In this experiment the total abundance of a community at the end of a transfer cycle did not considerably vary from transfer to transfer, meaning that we can assume that the total abundance at time *T* within a transfer cycle remained the same for all transfer cycles *k*,

(*N* ^(*k−*1)^(*T*) ≈*N* ^(*k*)^(*T*) ≡ *N* ^*∗*^(*T*)). Using this result, we divide the abundance of each ASV by the total abundance in a community to obtain relative abundances 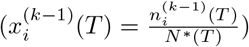, obtaining the following prediction for the mean abundance at the start of a transfer cycle

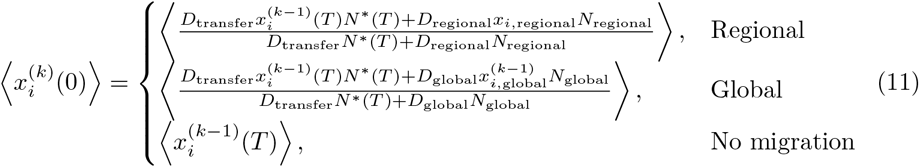

The impact that migration manipulations have on the initial relative abundance of the AFD and how they compare to the typical view of migration as a process that occurs at a constant rate is illustrated in Fig. S3. Parameter values are calculated using experimental details and can be found in Table 1 and simulation details of Eq. 9 can be found in S6 Text.

**Table 1.**
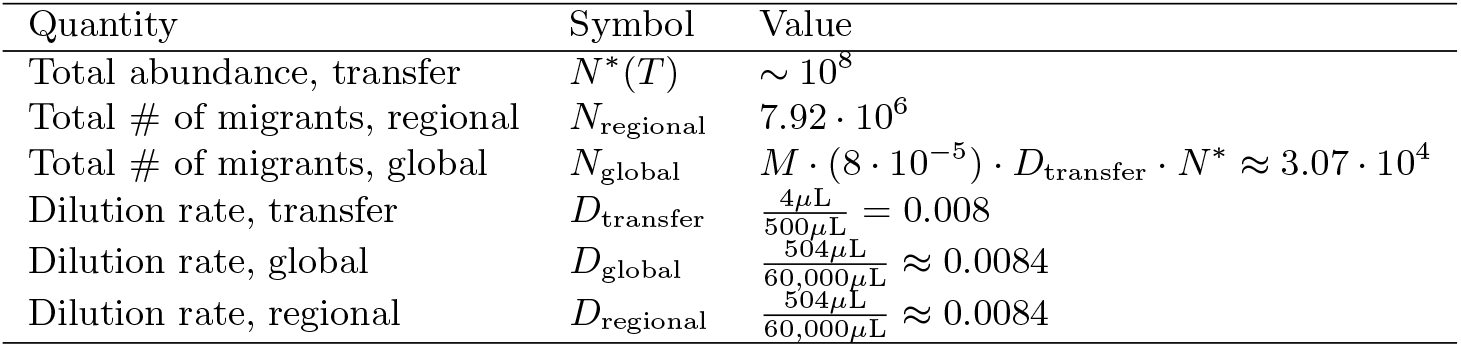
Experimentally imposed parameters that were used in this study [1]. Estimates of total community size were previously obtained. The calculation of *N*_global_ is the result the procedure used to pool samples from replicate communities into a single global pool at the end of each transfer, which was then used to inoculate communities at the start of the subsequent transfer.

### Treatment-specific migration statistics

We identified appropriate statistics to capture macroecological outcomes specific to the regional and global migration treatments. Significance was established in all instances using null distributions of statistics obtained via permutation.

#### Regional migration statistics

We first predicted that the mean relative abundance of an ASV in the assembled communities (⟨*x*_*i*_(*t*)⟩) should be affected by its relative abundance in the progenitor community (*x*_*i*,progenitor_) due to a mismatch in abundances. This prediction is justified, as correlation was weak between these two quantities (Fig. S4). Based on this observation, we sought to identify 1) how the correlation in ⟨*x*_*i*_(*t*)⟩ between no migration and migration treatments increased after the cessation of migration and 2) how the dependency of ⟨*x*_*i*_(*t*)⟩ on *x*_*i*,progenitor_ dissipated once migration manipulations ceased. The change in correlation coefficients (*ρ*) between transfers 12 and 18 was assessed using Fisher’s *Z* statistic [66]

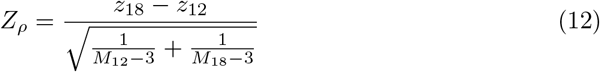

where 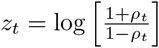 and *M*_*k*_ is the number of replicate communities at transfer *k*. The denominator represents the standard error of the numerator. Dependency on the progenitor was evaluated by fitting a regression between *x*_*i*,progenitor_ and the ratio of mean abundances in the regional and no migration treatments 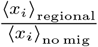. Regression fits were obtained at transfers 12 and 18 and the statistical significance of the change in slope was evaluated using a permutation-based *t*-test.

#### Global migration statistics

For the global migration treatment we predicted that fluctuations in abundance would decrease under migration among assembled communities. This prediction can be justified based on experimental details and the properties of statistical moments of distributions. To perform the global migration treatment, aliquots were taken of each replicate community on the *k* − 1th transfer cycle after *T* hours, pooled together and intermixed, then redistributed among the same communities at the start of the *k*th transfer cycle (Fig. 3). Ignoring fluctuations driven by sampling, the relative abundance of an ASV in the intermixed pool before it is redistributed will be 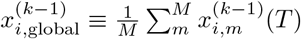, where the *M* ^*−*1^ prefactor accounts for an equal volume of each aliquot having been sampled. From this definition, we obtain the expected relative abundance in the global migration pool

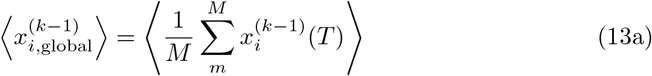

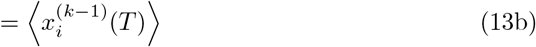

and the variance of relative abundance in the global migration pool

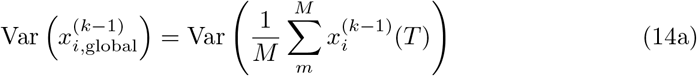

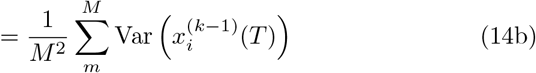

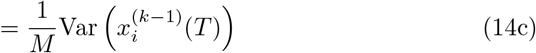

When an aliquot of the intermixed global migration pool is sampled and added to a community at the start of transfer cycle *k*, we find that the mean relative abundance at the start of a transfer cycle remains unchanged

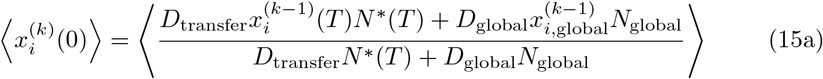

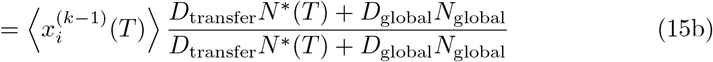

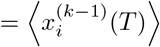

In contrast, when deriving the variance as fluctuations around the expected relative abundance, we obtain the following inequality

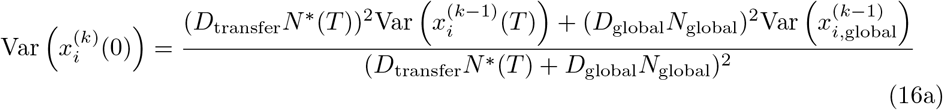

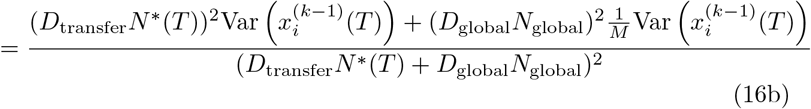

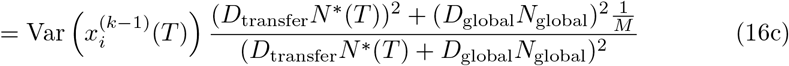

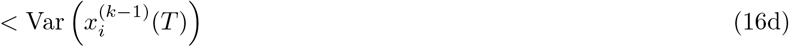

Therefore, under the SLM we expect global migration to alter the CV of relative abundances while leaving the mean unchanged.

We performed two types of analyses to test our fluctuation predictions: examining whether 1) the correlation in 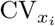 between no migration and global migration treatments and 2) the CV of the log-ratio of relative abundances between consecutive transfers increased after the cessation of migration. The log-ratio of relative abundances between two timepoints is defined as 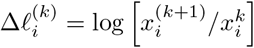. We elected to use 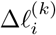 because 1) it can be interpreted as a discretized form of the per-capita growth rate and 2) as a ratio it cancels out any potential multiplicative time-independent sample biases [67, 68]. We first calculated the CV of 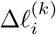 across communities at each transfer for each ASV, 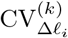. To contrast this measure of fluctuations *over replicates* with a measure of fluctuations over time *within a replicate*, we calculated the CV before and after the cessation of migration *for each replicate*, 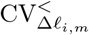 and 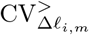. The effect of the cessation of migration on a per-replicate basis was examined using an *F* statistic designed to compare two CVs [69]

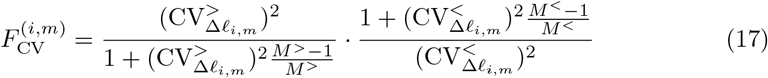

where *M*^*<*^ and *M*^*>*^ represent the number of transfers before and after the cessation of migration, respectively.

### Evaluating SLM predictions

Using our experiment-informed simulation, we determined whether shifts in macroecological quantitities due to migration could be explained by the SLM (S6 Text). To briefly summarize, we first simulated the experiment for 10^4^ uniformly drawn sets of {*τ, σ*}. We used Approximate Bayesian Computation (ABC) to identify the parameters where the simulated macroecological pattern had the lowest Euclidean distance with the observed pattern. We then used this optimal set of parameters to generate distributions of the macroecological pattern or summary statistic using 10^3^ SLM simulations. To evaluate whether migration was capable of altering a pattern under the SLM, we repeated our simulations across a grid of {*τ, σ*} combinations.

## Results

Using a high-replication community assembly experiment, we first examine whether universal macroecological patterns observed in natural communities could be recapitulated in a laboratory setting. We then examine whether the patterns we observed were susceptible to experimental manipulation. We rationalize our observations by extending the SLM to incorporate experimental details, allowing us to balance the effectiveness of the SLM as a minimal model of ecology with experimental realism. Using simulations, we obtain quantitative predictions and identify whether the macroecological consequences of a given form of migration could be captured by the SLM, establishing its capacity to predict patterns of microbial biodiversity.

### Macroecological patterns emerge in experimental systems

Predicting the outcomes of experimental manipulation is a vital goal of microbial ecology. With this goal in mind, we first determined the degree to which empirical macroecological patterns documented in observational data held in an experimental system (Figure 1a). The number of potential patterns one can examine is potentially very large. From previous works on natural microbial communities, we know that three main patters are sufficient to recapitulate several others [33]: 1) the mean and variance of species abundances displaying non-independence (Taylor’s Law), 2) the Abundance Fluctuation Distribution across independent sites (AFD) following a gamma distribution, and 3) the Mean Abundance Distribution across independent sites (MAD) following a lognormal distribution. Other commonly studied macroecological patterns (e.g., the Species Abundance Distribution, abundance-occupancy relationship), can be obtained and predicted as a consequence of these three patterns. Furthermore, these three patterns can be rationalized and predicted by a mean-field model, the Stochastic Logistic Model of growth [33].

We first quantified the variation in abundance that emerged in experimental communities. The average relative abundance varied over four orders of magnitude, meaning that a considerable degree of community-level variation can be maintained in a laboratory setting. We found that the shape of the relationship between the mean and variance of relative abundance across ASVs did not qualitatively vary across treatments, implying that Taylor’s Law can be applied [30]. Figure 1b shows that the variance of relative abundance was proportional to the square of the mean, corresponding to Taylor’s Law with an exponent equal to two. This value of the exponent implies that the coefficient of variation of relative abundances (CV) remained constant across ASVs. Fitting the exponent to each treatment for each transfer, we found a mean exponent of 2.1 ± 0.06, suggesting that despite the variation in typical abundance, the CV of relative abundances remained roughly constant across ASVs.

Given the existence of substantial fluctuations of abundance across replicate communities, we focused on the full distribution of abundances, known as the Abundance Fluctuation Distribution (AFD). To facilitate comparisons across ASVs and treatments, we rescaled the logarithm of the AFD for each ASV by its mean and variance (i.e., the standard score). We found that rescaled AFDs from different treatments tended to collapse on a single curve, implying that despite differences in experimental details, the general shape of the AFD remained invariant. Figure 2a shows that the bulk of the distributions generally followed the gamma distribution as observed in empirical data and predicted by the SLM (Eq. 2).

**Figure 2.**
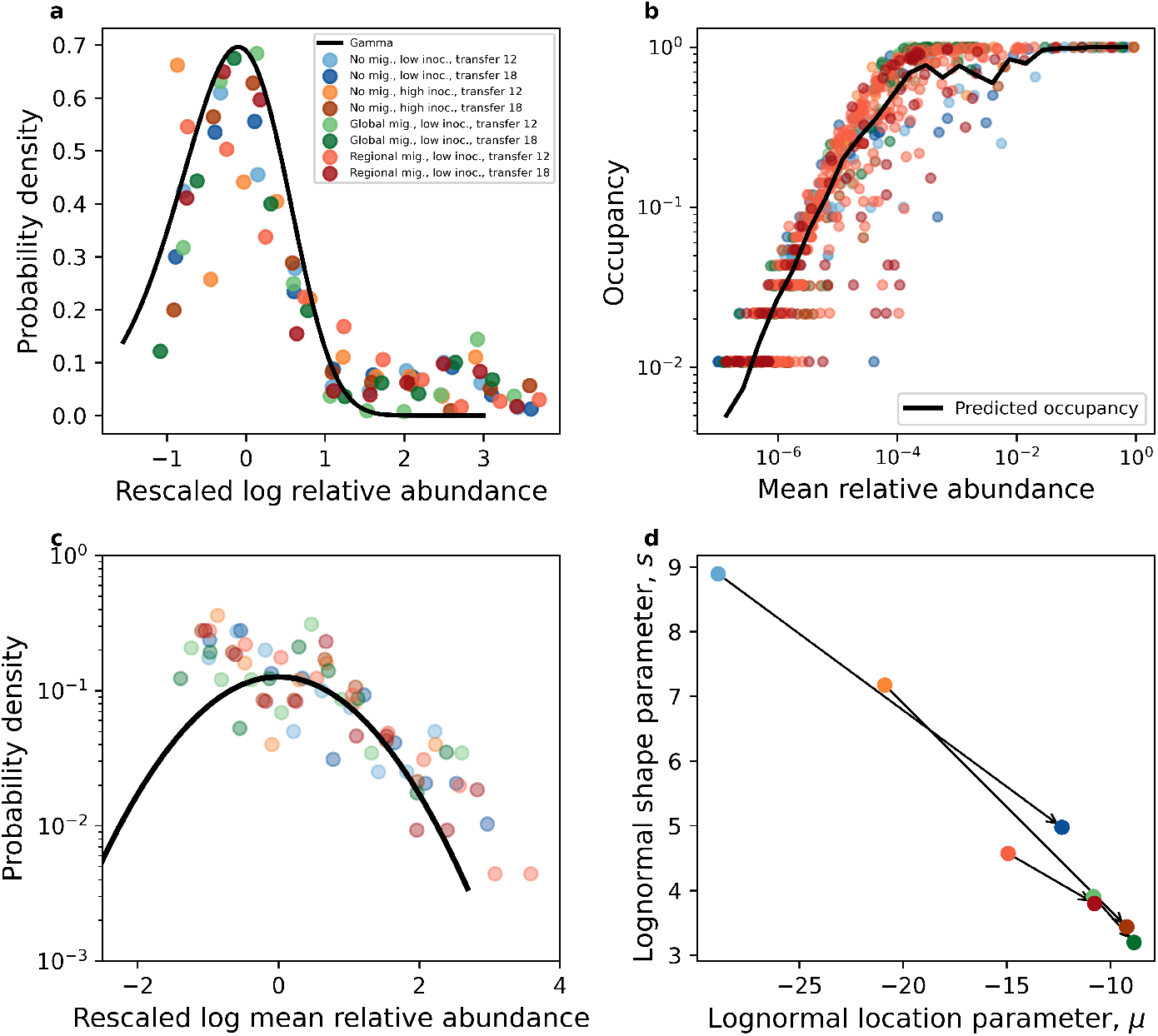
Macroecological patterns hold in experimental communities. Empirical macroecological patterns that were previously identified in natural microbial systems consistently arise in experimental communities [33]. **a)** The Abundance Fluctuation Distribution (AFD) tends to follow a gamma distribution across treatments. **b)** A gamma distribution that explicitly considers sampling successfully predicts the fraction of communities where an ASV is present (i.e., its occupancy). The prediction of the gamma (black line) was obtained by averaging over ASVs within a given mean relative abundance bin. **c)** The distributions of mean abundances are similar across treatments and largely follow a lognormal distribution and **d)** after the cessation of migration manipulations the two free parameters of the lognormal converged across treatments.

**Figure 3.**
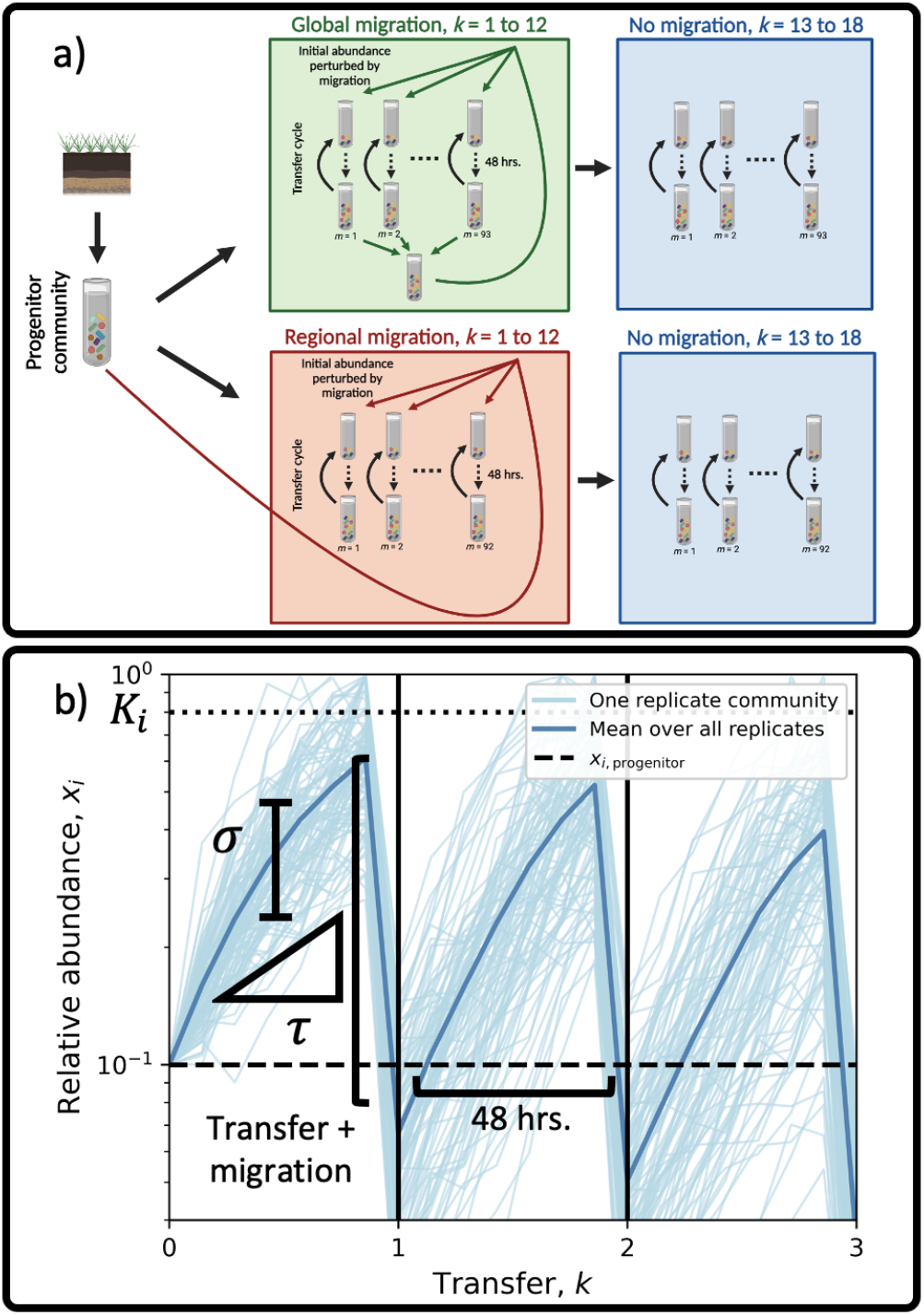
Incorporating experimental details into the Stochastic Logistic Model. **a)** Replicate communities were initiated from a single progenitor community that was isolated from soil. Communities were grown in microcosms containing a single carbon source for 48 hours, where they were then transferred to a microcosm with replenished resourced. This process constitutes a single transfer cycle. Migration was manipulated by altering the abundances of ASVs at the start of a transfer cycle. Two forms of migration were performed: regional and global. Regional migration represents a form of island-mainland migration, where aliquots of the progenitor community were added at the start of a transfer cycle. Global migration was manipulated by mixing aliquots of each replicate community at the end of a transfer cycle, which was then redistributed at the start of the subsequent transfer cycle. In both cases migration manipulations were performed for the first 12 transfers (1 to 12) and ceased for the remaining six transfers (13 to 18). Community assembly experiments were also performed with no migration manipulations for the entirety of the 18 transfers (not pictured). **b)** We modeled the ecological dynamics of each ASV within a given transfer cycle by incorporating the forms of migration performed in the experiment into the Stochastic Logistic Model of growth (Eq. S19). Under the SLM, the parameter *σ* controls the strength of environmental noise, represented here as the variation in trajectories within a transfer cycle. The parameter *τ* represents the minimum timescale of growth (inverse of the maximum rate of growth), controlling the rate that the population approaches its carrying capacity. Migration alters the relative abundance of an ASV at the start of a transfer cycle as a perturbation of initial conditions (Eq. 11). This experimental detail reduces the relative abundance of a given ASV if the relative abundance in a migration inoculum was lower than the carrying capacity (*K*_*i*_), where the ASV then proceeds to increase in abundance at a rate set by the timescale of growth *τ*. Conceptual diagram **a** was modified from Estrela et al. [1].

Abundance fluctuations offer only a partial view of community variation since patterns of presence/absence of community members could display non-trivial patterns. We therefore studied the fraction of communities where a given ASV was present, a measure known as occupancy. Given that the process of sampling ASVs within a community can be modeled as a Poisson process, the distribution of read counts can be derived from a gamma AFD which can be used to obtain occupancy predictions (Eq. 5). We find that the predictions of the SLM generally hold across treatments, with slight deviations at high values of observed occupancies (Fig. S1a). This trend is reflected in the distribution of relative errors in our occupancy predictions, where certain treatments appeared to have higher error values than others (Fig. S1b). However, upon examining ASVs that were present in global, regional, and no migration communities, we found that the error was not affected by migration (Fig. S1c). A permutation-based test was performed to establish the statistical significance of this observation. We found a significant, if slight, effect of migration, where it reduced the error of our predictions for both regional 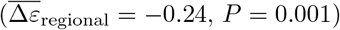 and global migration 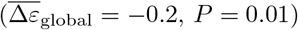. Regardless, Eq. 5 predicts the occupancy of the typical ASV across treatments with a high level of accuracy.

Leveraging this result, we investigate the relationship between the mean relative abundance across replicates and the occupancy of an ASV, more commonly known as the abundance-occupancy relationship [14, 63, 64]. Fig. 2b shows that all treatments follow the predictions of the gamma distribution. The small deviations from the prediction are likely driven by a combination of 1) averaging over a finite number of ASVs and 2) variation of *β*_*i*_ across ASVs [37]. The existence of a relationship between average abundance and occupancy in experimental communities is particularly striking, as it implies that the probability of observing a given community member is primarily determined by sampling effort (i.e., # reads) and mean relative abundance, despite differences in experimental details.

The success of the AFD in predicting occupancy and the observation that Taylor’s Law holds implies that differences in abundances are primarily driven by differences in their mean relative abundance. The distribution of the mean relative abundance across taxa (known as Mean Abundance Distribution, MAD) in observational data is well described by a lognormal distribution ( [33]; Eq. 8). This observation can be seen as an across-community extension of the observation that the distribution of abundances within a single microbial community can be captured by a lognormal distribution [12]. The low richness of experimental communities ( ∼ 10 ASVs per community) makes it difficult to study the MAD of a given treatment, requiring ASVs to be pooled across treatments and transfers. Given the low sampling level, the resulting empirical MAD can be captured by a lognormal (Fig. 2c).

It is important to note that the lognormality of the MAD itself does not validate or invalidate the SLM. Rather, under the SLM the observation that the mean relative abundance and variance are not independent implies that the mean is proportional to the carrying capacity 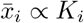, so evidence of a lognormal MAD informs us of the distribution of parameters used in the SLM, not the SLM itself.

In order to assess the effect of migration on the average abundances, we compare the MAD between transfer 12 (end of migration treatment) and transfer 18 (the last transfer in the experiment). Fig. 2d shows that the two MADs have a similar shape, suggesting that the MAD as a macroecological pattern consistently converged to a similar form, irrespective of the migration treatment.

### Testing the macroecological effects of migration

Our analysis shows that demographic manipulations of experimental communities do not induce *qualitative* changes in macroecological patterns (e.g., altering the form of Taylor’s Law). This result is consistent with the observation that natural microbial communities display similar patterns [22, 33]. However, the *quantitative* effects of these experimental manipulations is unclear. To determine whether these effects exist and the degree that they can be predicted, we examined deviations in macroecological quantities, incorporated experiment-specific forms of migration into the SLM, and tested their predictive capacity.

It is clear from the previous section that the SLM is a useful descriptive model of experimental microbial communities. Therefore, it is reasonable to expect that a form of the SLM that incorporates migration could serve as a predictive model. While a constant rate of migration appears intuitive and can be incorporated into the SLM (S4 Text), this model does not reflect the details of the experiment. Rather, in this experiment migrants only entered communities at the start of a given transfer cycle (Fig. 3a). This detail corresponds to a model where the effect of migration can be modeled as an experimentally-induced perturbation on the initial abundance of an ASV within the *k*th transfer cycle 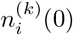; Materials and Methods, S6 Text). The full time-dependent solution of the SLM which depends on initial conditions has previously been solved [65], allowing one to model the temporal evolution of the AFD in response to experimental perturbations that alter initial abundances and compare their effects to AFDs with a constant rate of migration (S5 Text; Fig. S2).

While a time-dependent SLM appears appropriate, additional decisions must be made about how parameters are determined and the number of variables (i.e., community richness). Using rarefaction curves and previously established richness estimation procedures [70], we found that the richness of the progenitor community was ∼ 100 fold greater than the richness of a typical assembled community (Fig. S3). Migration has little effect on richness, as estimates are only slightly higher among communities that experienced migration while remaining orders of magnitude lower than the richness of the progenitor community. This result suggests that the carrying capacity of a given ASV played a primary role in determining its survival, raising the question of how to 1) specify the carrying capacity of an ASV in the assembled communities (*K*_*i*_) and 2) determine how *K*_*i*_ relates to the abundance of an ASV in the progenitor community. The observation that Taylor’s Law holds in the no migration treatments implies that the coefficient of variation of growth fluctuations was constant across ASVs (i.e., *σ*_*i*_ ≈ *σ*), meaning that the mean relative abundance of an ASV across communities is proportional to its carrying capacity 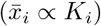. Given that 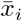 follows a lognormal distribution, *K*_*i*_ should also follow a lognormal.

The weak correlations between the mean relative abundance after the cessation of migration 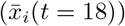 and the relative abundance observed in the progenitor (*x*_*i*,progenitor_) for all treatments supports the view that the *K*_*i*_ of a given ASV in the assembled communities is independent of its abundance in the progenitor community (Fig. S4). However, this observation suffers from survivorship bias, since we can only compare values of 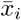 and *x*_*i*,progenitor_ for ASVs that were actually present in both the progenitor and assembled communities (i.e., ASVs with *K*_*i*_ *>* 0). We found that distributions of *x*_*i*,progenitor_ differed depending on whether a given ASV was present in the assembled communities, with ASVs that were present in assembled communities having a higher relative abundance in the progenitor (Fig. S5a). A permutational Kolmogorov–Smirnov test found that this shift was significant, a result that held for all experimental treatments (Fig. S6). We modeled the dependency between the occupancy of an ASV is non-zero and its abundance in the progenitor using logistic regression (Fig. S5b). This statistical dependence between the progenitor and assembled communities, along with details related to sampling (i.e., number of reads, Fig. S7), were incorporated into our numerical simulations of the SLM (Materials and Methods, S6 Text). Information about the statistical inference of the two free SLM parameters can be found in S7 Text.

### Experiment-agnostic macroecological patterns

We first examined the quantitative effect of migration using macroecological patterns that have been unified by the SLM: the AFD and Taylor’s Law. Fig. 4a-c shows that the AFDs of ASVs rescaled by their mean and variance differ between transfers 12 (migration present) and 18 (migration absent). This shift is particularly notable in the regional migration treatment, whereas the global migration treatment AFDs appear to be the most similar.

**Figure 4.**
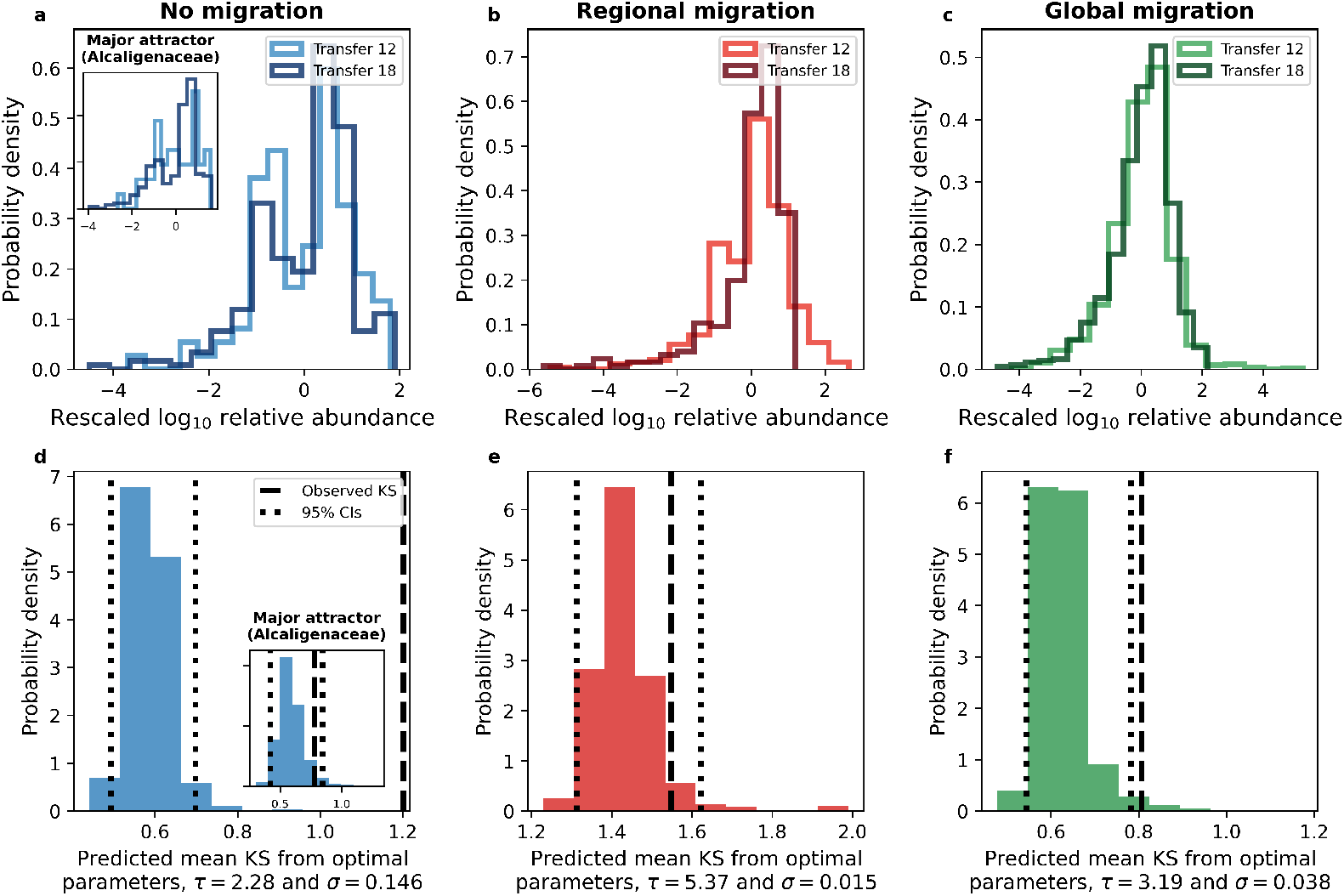
Migration impacts the shape of AFDs. **a-c)** By rescaling log_10_ transformed AFDs for each ASV, we can examine how the shape of the AFD changes before and after the cessation of migration. We quantified this shift by estimating the KS distance between ASVs before (transfer 12) and after (transfer 18) the cessation of the migration treatment for each ASV, then taking the mean over ASVs. **d-f)** Using optimal parameter combinations identified by ABC, we found that the SLM predicted reasonable KS statistics for regional migration as well as within the major attractor of the no migration treatment (inset of sub-plots **a**,**d**), with borderline successful predictions for the global migration treatment.

To examine the shift in the AFDs we calculated the KS distance between AFDs from transfer 18 and 12 for each ASV, focusing on ASVs present in every replicate community, and then calculated the mean KS over ASVs. Using Approximate Bayesian Computation (ABC), we identified the {*τ, σ*} pair that best explained the observed mean KS statistic (Materials and Methods; Text S6). We then used the optimal set of parameters to generate distributions of mean KS using 10^3^ SLM simulations. This significant discrepancy can be explained by the existence of multiple attractors, a scenario where certain ASVs had high abundances in a subset of communities and low abundances in others, a qualitative deviation from the SLM [1]. Focusing on no migration communities belonging to the major attractor (∼70% of communities, Table S2), we found that the SLM could predict the observed mean KS (insert of Fig. 4d). The effect of regional migration was also captured by the SLM (Fig. 4e), with the observed statistic for global migration communities lying slightly outside predicted range (Fig. 4f). However, manipulating the SLM parameters revealed that the AFD exhibited the largest systematic deviation via migration for the regional migration treatment (Fig. S8). This result suggests that the inability of the SLM to fully capture the change in AFDs for the global migration treatment was due to its uninformative nature for this particular form of migration.

We then analyzed how migration impacted the exponent of Taylor’s Law (Fig. 5a-f). Under regional micgration, the exponent is lower than that of the no migration at transfer 12 (Fig. 5a,b), but approaches the result of the no migration treatment by transfer 18 (Fig. 5d,e). Similarly, the intercept of regional migration treatment is initially higher but then approaches the value of the no migration treatment at treatment 18, as seen by the range of values on the y-axis. In contrast, there is little change in the values of the exponent or intercept for the global migration treatment. In line with our predictions, there is no significant change in the exponent (*t*_exponent_ = −1.1, *P* = 0.2) or the intercept (*t*_intercept_ = −0.6, *P* = 0.6) between transfers 12 and 18 for the communities that did not undergo migration. We found that the global migration did not alter the exponent (*t*_exponent_ = −0.4, *P* = 0.7) or the intercept (*t*_intercept_ = −0.4, *P* = 0.7). Contrastingly, regional migration significantly altered both the exponent (*t*_exponent_ = 3.3, *P* = 0.02) and the intercept (*t*_intercept_ = 2.75, *P* = 0.03).

**Figure 5.**
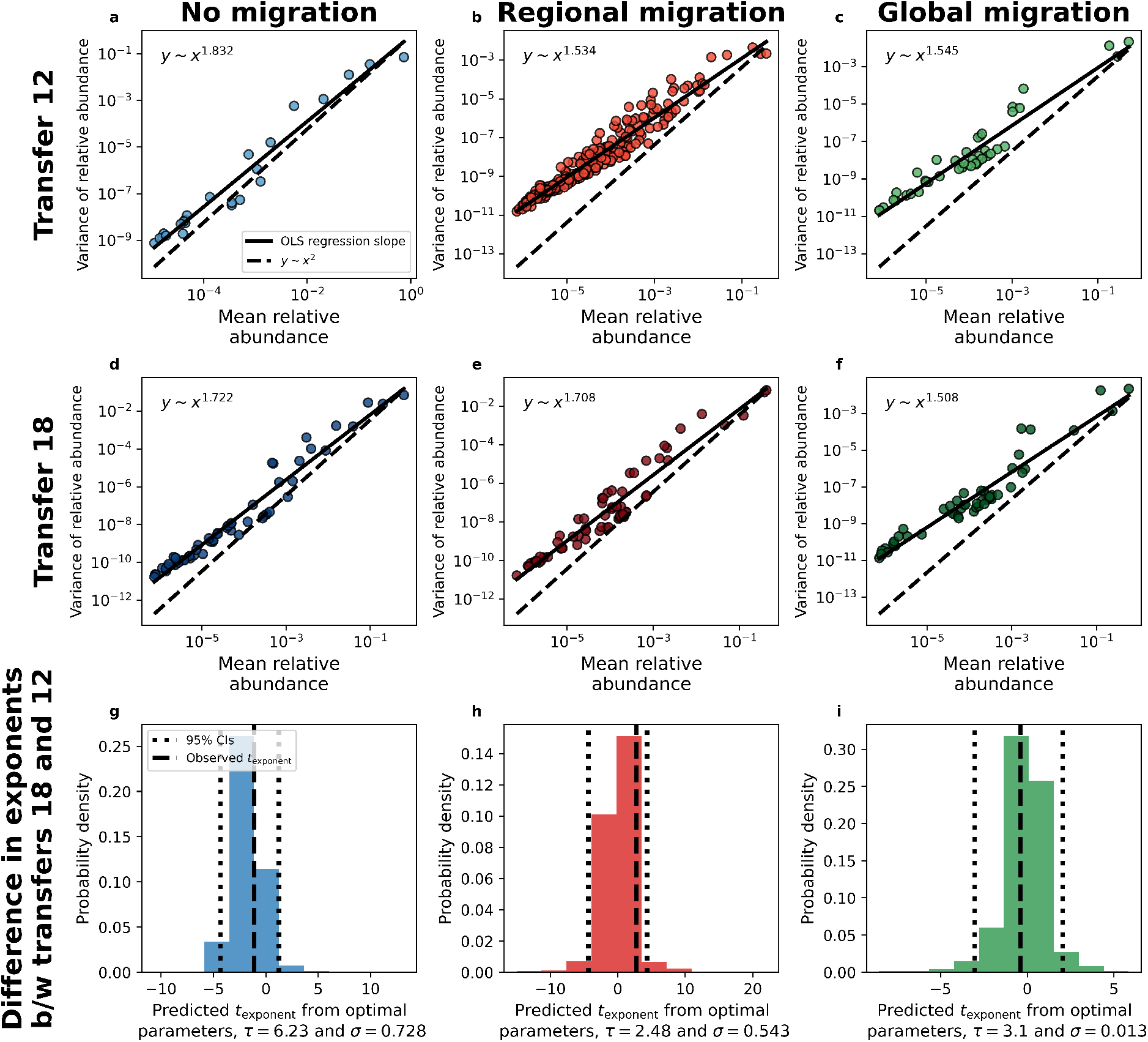
Regional migration impacts the exponent of Taylor’s Law. **a-f)** By examining the exponent of Taylor’s Law for each treatment we found that the exponent only considerably changed after the cessation of migration for the regional migration treatment. Each data point represents statistical moments calculated across replicate communities for a single ASV from a single experimental treatment. **g-i)** Using our ABC estimation procedure, the SLM succeeds in predicting this increase in the exponent for regional migration.

By repeating the same ABC procedure outlined for the AFD we found that the distribution of exponents generated for an optimal set of parameters could predict the observed values of *t*_exponent_ across all migration treatments (Fig. 5d-f). However, by examining *t*_exponent_ and *t*_intercept_ across a grid of parameter combinations we found that the statistic was again only informative for the regional migration treatment (Figs. S9, S10). Our results demonstrate that the AFD and Taylor’s Law were primarily informative of the effects of regional (island-mainland) migration.

### Experiment-specific macroecological patterns

The macroecological patterns we have examined up to this point can be explained by the SLM, though the AFD was not substantially altered by migration. It is useful to supplement our analyses by considering novel macroecological patterns that are likely to be altered by migration. Given the details of the experiment, expected regional migration to alter typical abundance, whereas global migration would alter fluctuations in abundance (Materials and Methods).

#### Regional migration

Given the difference in abundance between assembled communities and the progenitor, we predicted that regional migration would alter the mean abundance of an ASV. We examined paired MADs for the regional and no migration treatments *before* (*k* = 12) and *after* (*k* = 18) the cessation of migration. The correlation between MADs was initially low and non-significant at transfer 12 but had significantly increased by transfer 18 (Fig. 6a,b; *Z*_*ρ*_ = 2.7, *P* = 0.01) [66]. Our SLM simulations recapitulate this observation, as we are able to predict the observed value for the optimal parameter combination of *σ* and *τ* from ABC (Fig. 6c) and over a specific range of parameter combinations (Fig. S11a,b). These results are consistent with the interpretation that ASVs revert to their previous typical abundance (captured by the carrying capacity) once island-mainland migration ceased.

**Figure 6.**
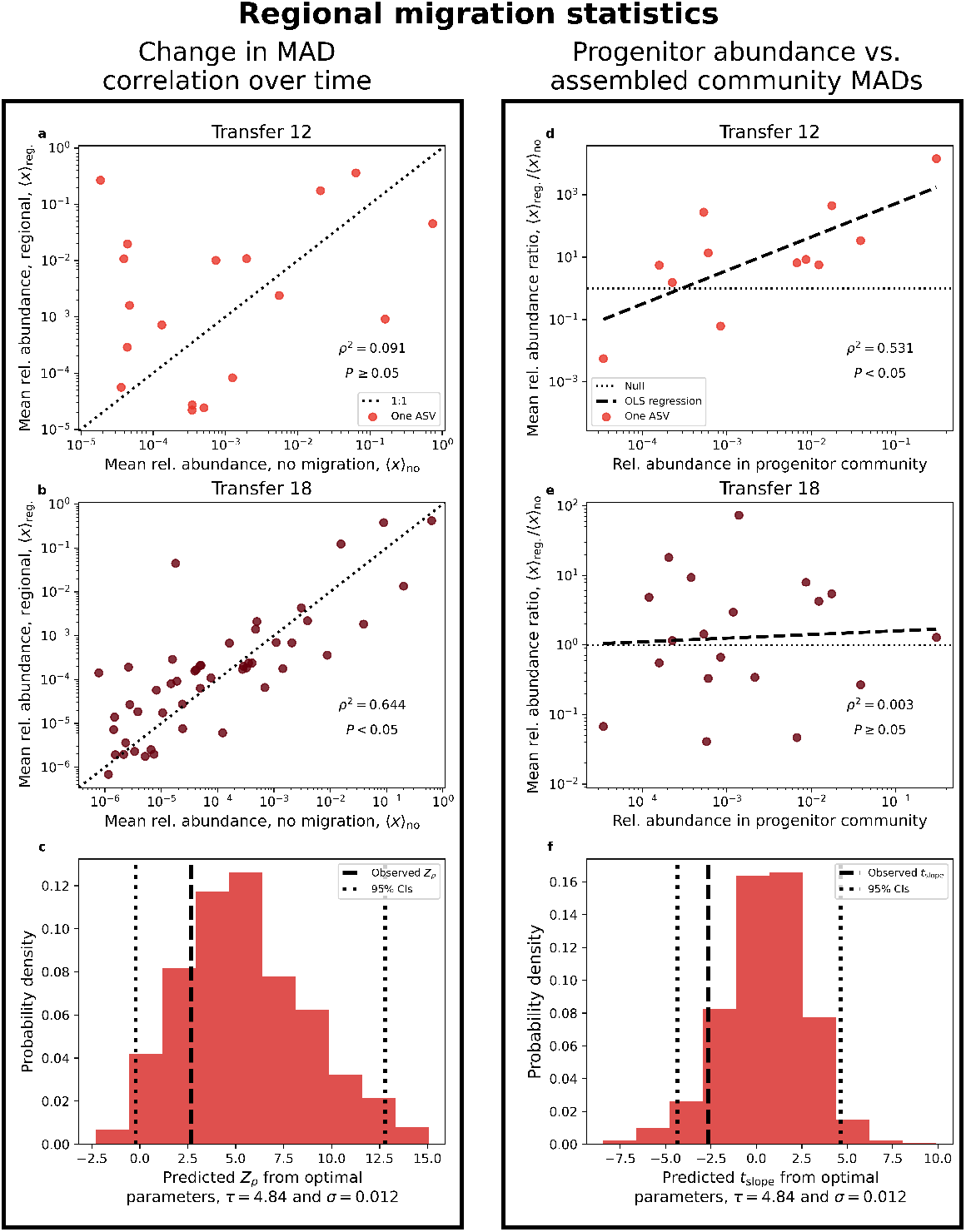
Regional migration alters typical abundance in a manner consistent with SLM simulations. The properties of the MAD were examined to evaluate the impact of migration over time. **a)** At transfer 12, the last transfer with migration, we found no apparent correlation between the MADs of the regional and no migration treatments. **b)** Contrastingly, the strength of the correlation rapidly increased by transfer 18. The significance of this difference can be evaluated by calculating Fisher’s *Z*-statistic, a statistic that can be applied to correlations calculated from simulated data. **c)** By performing simulations over a range of *σ* and *τ*, we can visualize how the observed value of *Z*_*ρ*_ compares to simulated distributions and identify reasonable parameter regimes **d)** as well as the relative error of our predictions. **e)** The ratio of the MAD between regional and no migration provides a single variable that can be compared to the abundance of an ASV in the progenitor community. We found that there was a positive significant slope at transfer 12 that dissipated by transfer 18, reflecting the cessation of migration. Only sub-plots **c** and **f** contain simulated data.

We then sought to determine whether the effect of regional migration depended on the abundance of an ASV in the progenitor community. To examine this dependence, we studied the relationship between the ratio of the mean abundance of an ASV in the regional and no migration treatments 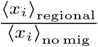 and its abundance in the progenitor community. We found that at transfer 12 there was a strong relationship between the two quantities (statistically significant via permutation). By transfer 18 the slope is statistically indistiguishable from zero (Fig. 6d,e). The observed change in slopes is significant (*t* = −2.6, *P* = 0.03) and can be reproduced using the SLM for certain *σ, τ* parameter combinations. These combinations overlapped with those that reproduced observed estimates of *Z*_*ρ*_ (Fig. 6f; Fig. S11c,d), providing further validation of the SLM.

#### Global migration

Accourding to the prediction of the model, global migration would strictly alter fluctuations in ASV abundance across replicate communities while leaving the MAD unchanged (Materials and Methods). Specifically, the correlation in the MAD between global and no migration treatments should remain unchanged before and after the cessation of migration manipulations. This prediction holds, as there is no significant change in the correlation between transfers 12 and 18 (Fig. S12; *Z*_*ρ*_ = 0.2, *P* = 0.4). However, there was also no significant increase in the strength of correlation for the distribution of CVs (*Z*_*ρ*_ = 0.3, *P* = 0.6), meaning that the cessation of migration did not considerably alter fluctuations in abundance.

It is possible that two time points are insufficient to detect the impact of global migration. Detecting changes in fluctuations often requires more observations than detecting changes in typical values. As a solution, we leverage the higher sampling resolution of the global migration treatment to examine the change in abundance between time points (Δ*ℓ*), as a subset of global migration communities were sequenced at each of the 18 transfers (Table S1). The mean change in abundance across replicate communities at transfer *k* (⟨Δ*ℓ*^(*k*)^⟩) tends to relax towards a value of zero around the sixth transfer and remained there throughout the rest of the experiment (Fig. S14). This trend indicates that the abundances of ASVs reached stationarity with respect to the start of the experiment by transfer 6, allowing us to examine equal intervals of transfer cycles before (7 − 12) and after (13 − 18) the cessation of migration.

Using permutational *t*-tests we determined whether ⟨Δ*ℓ* ^(*k*)^⟩ was altered after the cessation of migration, controlling for ASV identity. There was no evidence that ⟨Δ*ℓ*^(*k*)^⟩ changes in either the no migration treatment 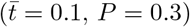 nor in the global migration treatment 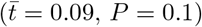. This result is consistent with that prediction that global migration would not alter typical abundances.

Turning to fluctuations, we examined the CV of Δ*ℓ* ^(*k*)^. As predicted, the distribution of 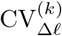 does not increase for the no migration treatment 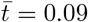 *P* = 0.8; Fig. 7a). In the global migration treatment, the coefficient of variation 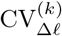 significantly increases after the cessation of migration (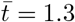, *P <* 10^*−*3^; Fig. 7b). This result is consistent with our prediction that global migration dampens fluctuations across communities. However, while our SLM simulations succeeds in predicting *t̄* for the no migration treatment, they fail to capture the effects of the global migration treatment (Fig. 7c,d).

**Figure 7.**
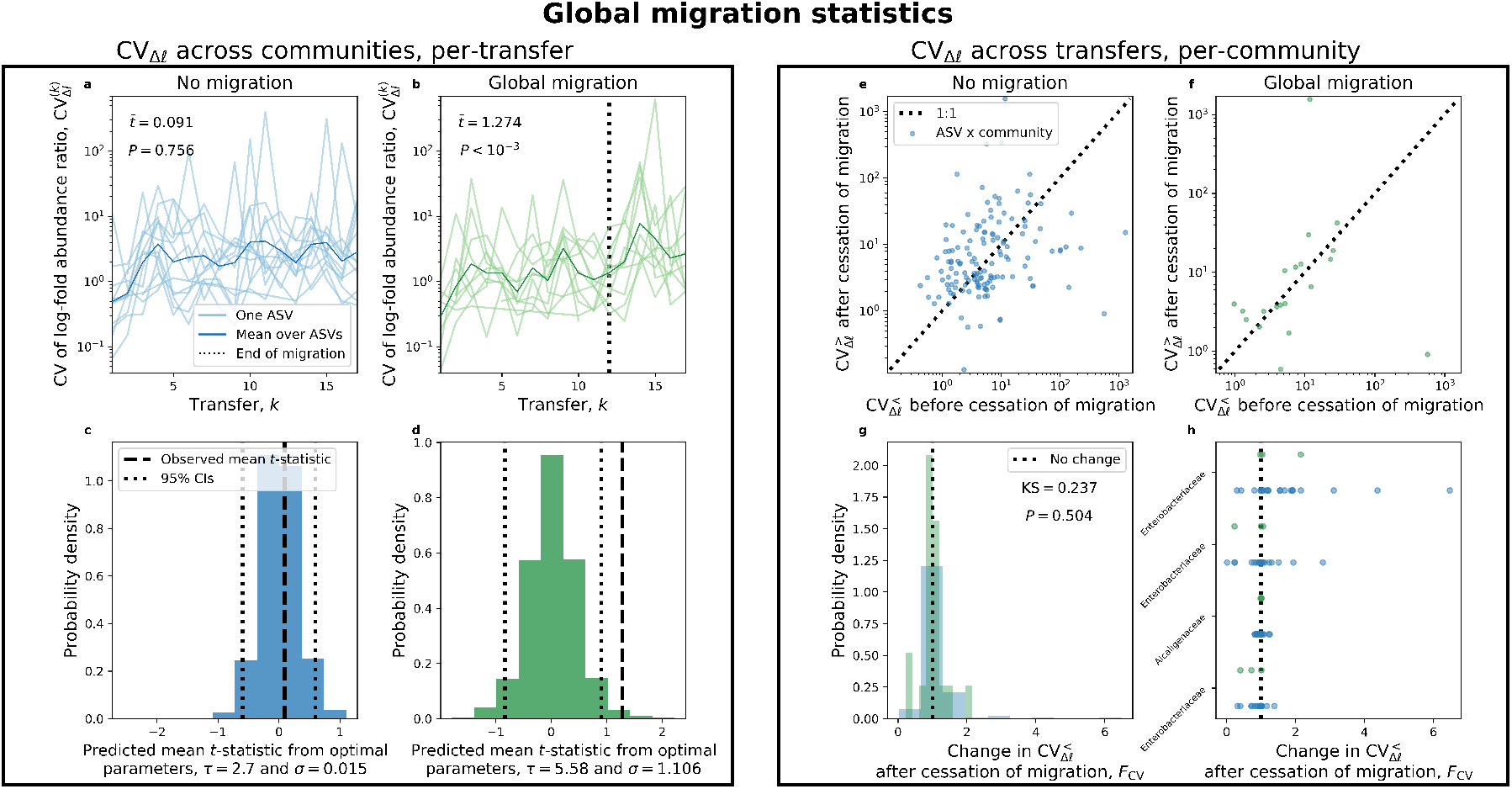
SLM simulations are unable to reproduce log-fold fluctuations *across* communities under global migration. By leveraging the entire timeseries we can examine how the coefficient of variation of Δ *ℓ* was altered by migration for two scenarios: 1) fluctuations across replicate communities at a given transfer 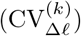 and 2) fluctuations before 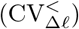 and after 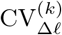 the cessation of migration within a given community. **a)** As expected, there is no change in 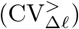 for the no migration treatment. **b)** However, 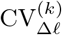 tends to slightly increase after transfer 12 for global migration, a shift that was found to be significant using a *t*-test where transfer labels were permuted for each ASV. **c, d)** Our SLM predictions succeeded for the no migration treatment, but was unable to capture observed values of 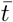 under global migration. These results pertaining to the fluctuations across communities can be contrasted with fluctuations within a community. **e, f)** We did not observed a systematic shift in the CV before and after the cessation of migration for either treatment. **g)** By calculating the difference between two CVs (*F*_CV_) for each ASV in each replicate community, we do not observe a significant difference between migration treatments using a KS test constrained on ASV identity. **h)** This conclusion is confirmed by investigating the change in the CV for high occupancy ASVs, as there is no systematic difference between migration treatments.

Experimental details may explain why the effect of the global migration treatment, while significant, was not particularly large, as the size of the inoculum was nearly two orders of magnitude smaller than that of regional migration (Table 1). However, this experimentally-imposed parameter does not explain why the observed value of 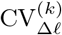 lay outside the predictions of the SLM. Attractor status also does not explain this discrepancy, as the global migration communities were previously classified as belonging to the same attractor (Table S2, [1]). Instead of evaluating fluctuations *across* communities, we focused on fluctuations over time *within* individual communities to evaluate the effect, if any, of global migration. We see that estimates of the CV before 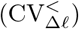 and after the cessation of migration 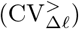 are similar for both no and global migration treatments (Fig. 7e,f). Using a measure of the change in CV, we find that distributions of both treatments highly overlap, meaning that the change in the CV *within a given community* does not considerably increase in the global migration treatment (Fig. 7g). Splitting said distribution into high occupancy ASVs reinforces this conclusion, as there is no systematic increase in the change in CV for the global migration treatment (Fig. 7h).

## Discussion

The main result of this study is the demonstration that experimental microbial communities grown on a single carbon source can sustain large ecological variation. This variation provides the means to assess distributions of typical abundances across several orders of magnitude, a prerequisite for examining broad probabilistic patterns of diversity and using a macroecological approach. Meeting this criterion allowed us to document the existence of macroecological patterns that have only been documented in naturally occurring communities, suggesting a quantitative equivalence between experimental and observational studies despite the controlled nature of artificially maintained communities. However, the repeatable maintenance of variation observed in the face of demographic manipulations was likely contingent on the high level of variation present in the progenitor community. This figurative raw material is analogous to the need for genetic variation to exist before selection can occur [71–73], the absence of which would preclude the possibility of macroecological investigations.

Characterizing robust empirical patterns in artificial communities is a key step toward identifying predictive ecological models. The existence of patterns predicted by the SLM in artificial communities provided an opportunity to evaluate the macroecological consequences of experimental manipulations. Our approach of extending the SLM, an empirically validated model of microbial community composition, using experimental details proved to be a useful framework for identifying treatments that were capable of generating macroecological effects. We examined the effects of two different forms of migration: regional (island-mainland) and global (fully interconnected metacommunity) [1]. As expected, regional migration altered macroecological patterns of typical relative abundance in a manner that was captured by the SLM using minimal free parameters. The results of our regional migration analyses demonstrate that the SLM can link experimental communities and macroecological patterns.

Regarding global migration, we predicted that the treatment would primarily alter ASV fluctuations around their typical abundance [59]. We observed no change in the abundance fluctuations of community members after the cessation of migration *within individual communities*. However, we found that variation *across communities* tended to increase after the cessation of migration. This trend is consistent with our hypothesis regarding the effect of global migration, but one that cannot be reproduced by the SLM. This disagreement between fluctuations *across* and *within* communities suggests that experimental communities readily reach a state where their statistical properties do not change over transfer cycles, but can vary from community-to-community. In physics parlance, experimental communities are stationary, but may not necessarily be ergodic [74]. The existence of alternative stable-states are a likely explanation, as they are common in experimental microbial communities [56, 75, 76] and can be induced via fluctuations intrinsic to serial dilution setups [77]. In addition, their ergodicity-breaking effects have been used to explain invasion dynamics in experimental microbial communities [78]. The SLM can incorporate such states [37], though prior efforts required extensive temporal samples. Regardless, we can make two claims about the impact of alternative stable-states. First, the quantity of migrants chosen for the global migration treatment was sufficient to alter attractor status, but insufficient to alter fluctuations in abundance within a given community. Second, macroecological patterns captured by the SLM persisted despite the pervasiveness of alternative stable-states, implying that heterogeneous outcomes of community assembly do not qualitatively alter macroecological patterns.

In this study we focused on patterns relating to the typical abundance and fluctuations in abundance across communities as well as over time. Noticeably, we did not investigate patterns that likely require the explicit invocation of interactions between community members (e.g., the correlations of abundance fluctuations), as the addition of interaction terms into the SLM would require several assumptions about the network of interactions and their magnitude. However, the absence of interaction terms in the SLM does not mean that interactions did not contribute to the macroecological patterns we observed. Models such as the SLM are known as “mean-field” models, where interactions between community members are permitted, but their effects on the dynamics of a given community member are determined by the mean dynamics of all community members [79]. Specifically, the SLM as presented in this study will continue to hold if the effect of community interactions is restricted to the parameters of the SLM (e.g., carrying capacity). Investigating correlations in abundance between community members, which can be understood as an outcome of specific ecological interactions rather than the mean effect of the community, likely requires models that go beyond phenomenology by explicitly considering mechanisms such as resource consumption [13, 80]. Indeed, consideration of resource consumption has proven critical for investigating the evolutionary dynamics of microorganisms in an ecological context [23].

Recent developments on the predictability of community function (e.g., total biomass, polysaccharide hydrolysis, resource excretion, etc.) point towards new avenues of exploration for microbial macroecology. There is increasing evidence that the functional profiles of experimental communities tend to follow quantitative rules that are amenable to mathematical modeling [81–84]. Extending studies of microbial macroecology beyond patterns of abundance and community composition to the level of function would fully realize the physiological and energetic breadth that allowed macroecology vital to advance our understanding of macrobial life [5].

It is worth discussing how the experimental implementation of migration relates to natural environments. Microbial community assembly experiments often examine ecological dynamics as a “boom-and-bust” phenomenon, where abundances are initially low and proceed to reach a carrying capacity. This decision is often made due to the unwieldy nature of managing an array of continuous cultures (i.e., chemostats). However, this experimental design may reflect pulse-like forms of migration in nature, as boom-and-bust dynamics can be found across diverse ecosystems. Examples of environments where boom-and-bust dynamics occur include pitcher plants [85], particles of organic matter in the open ocean [86, 87], and even the human gut [88, 89]. Boom-and-bust dynamics can serve as a useful model for environments where resources are periodically supplied, as migrants must persist until the environment becomes favorable to growth (e.g., surviving in a metabolically inactive state [90–92]). Migration in such systems is difficult to quantify, though recent evidence suggests that it is sufficiently high such that microbes are rarely endemic to the environment in which they are found [93]. Appropriate models of migration may be necessary to investigate the macroecology of natural microbial communities.

Finally, considering how the results presented here shape the microbial view of macroecology. The discipline of macroecology was originally conceived as an explicitly non-experimental form of investigation [5]. Analysis of the origin and development of macroecology provides two historical explanations for the initial rejection of experimental approaches: 1) large-scale community-level experiments were often impractical and 2) producing generalities from experiments has proven to be difficult [94]. Our results demonstrate that these two constraints are ameliorated by the features of microbial communities, providing counter evidence to recent claims that statistical distributions do not provide information about ecological mechanisms [95]. The timescales, abundance, and comparative ease with which ensembles of communities can be maintained and manipulated make microorganisms an ideal system for testing quantitative macroecological predictions.

## Supporting information

experimental_macroecology_supplement

## Acknowledgments

We thank S. Estrela for her assistance in reprocessing the data. We thank S. Bubnovich, M. Dal Bello, A. Goyal and members of the qEcoEvo group at ICTP for helpful discussions. We thank B.H. Good, O. Mazzarisi, M. Sireci, and N. I. Wisnoski for their comments on the manuscript. This work was supported by the NSF Postdoctoral Research Fellowships in Biology Program under Grant No. 2010885 (W.R.S.). Á.S. acknowledges support from Grant PID2021-125478NA-I00 funded by MCIN/AEI/10.13039/501100011033 and by “ERDF A way of making Europe”. Conceptual diagrams made with BioRender.com.

## Author contributions

W.R.S., Á.S., and J.G. conceptualized the project, developed the mathematical models, and wrote the manuscript. W.R.S. performed all analyses.

## Data and code availability

All code written for this study is available on GitHub under a GNU General Public License: https://github.com/wrshoemaker/experimental_macroecology. Raw data was previously generated and can be accessed through the original manuscript [1]. Processed data for this project is available on Zenodo (DOI: 10.5281/zenodo.8393848).

