## Supplementary material for "Macroecological patterns in experimental microbial communities": experimental_macroecology_supplement

#### Supporting Information

### S1 Text: Additional experimental details

Experimental data was obtained from a previous study where a large number of replicate ecological communities originating from a single progenitor soil sample were propagated in controlled laboratory conditions under regional or global migration treatments (Table S1; [1]). A given replicate community was initiated by inoculating 4 $\mu$L (low inoculum) or 40 $\mu$L (high inoculum) of the source community into 500 $\mu$L of M9 minimal media with 0.2% glucose into a well of a 96 deep-well plate (VWR) at 30°C under static conditions. Transfers were performed every 48 hours, 18 times in total for all replicate communities with a dilution rate $D_{\text{transfer}} = 0.0084$ (Table 1).

The transfer procedure was modified to manipulate the effects of different forms of migration. For the regional migration treatment, a 4 $\mu$L aliquot of the soil supernatant was added to the 4 $\mu$L aliquot from the previous transfer. For the global migration treatment, 4 $\mu$L aliquots of all replicate populations were pooled, resuspended, and diluted 10,000-fold. A 4 $\mu$L aliquot of the diluted solution was added to the 4 $\mu$L aliquot from the previous transfer for each replicate community. At the end of each 48 h. transfer period samples from each replicate community were mixed with 40% glycerol, cryopreserved at -80°C, and DNA was extracted and sequenced.

DNA extraction was performed using a QIAGEN DNeasy 96 Blood and Tissue kit. Library preparation for 16S rRNA amplicon sequencing of the V4 region was performed as previously described [2] and PCR products were purified and normalized using the SequalPrep PCR kit (Invitrogen). Sequencing was performed on an Illumina MiSeq (2x250 bp paired-end) and raw reads were processed for demultiplexing and barcode, index, and primer removal using QIIME v1.9 [3]. The number of communities sequenced at each transfer for each treatment can be found in Table S1. In this study we reprocessed all raw FASTQ data from the original study to obtain Amplicon Sequence Variants (ASVs) using DADA2 [4]. We reprocessed the data using the pooled option inference option so that ASVs with an abundance of one in a given community (i.e., singletons) could be inferred, allowing us to examine the entirety of the empirical sampling distribution. The attractor status of a given replicate community was assigned as previously described [1]. Additional detail can be found in the study that originally presented the experiment [1].

#### S2 Text: Error estimates

Error between observed and predicted quantities was estimated as the relative error.

$$\varepsilon = \left| \frac{\text{Obs.} - \text{Pred.}}{\text{Obs.}} \right| \tag{S1}$$

The difference in relative error between two treatments was estimated as

$$\overline{\Delta\varepsilon} = \frac{1}{S_{\text{obs}}} \sum_{i=1}^{S_{\text{obs}}} \log \left[ \frac{\varepsilon_{\text{mig.}}}{\varepsilon_{\text{no mig.}}} \right] \tag{S2}$$

### S3 Text: Phenomenological logistic growth from a consumer-resource model

We provide a heuristic explanation for how a deterministic phenomenological model of logistic growth can serve as a useful model for a population in a closed system with an initial concentration of supplied resources (i.e., a given transfer cycle illustrated by Fig. 3). We start with a system of equations where one community member of abundance $x$ consumes a single resource $c$ to grow at rate $r(c)$

$$\frac{dx}{dt} = r(c)x \tag{S3a}$$

$$\frac{dc}{dt} = -\frac{r(c)x}{Y} \tag{S3b}$$

where $Y$ represents the cell yield per-unit resource and the growth rate can be represented as Monod kinetics $r(c) = r_{\max}\frac{c}{c+K_c}$ where $r_{\max}$ is the maximum possible rate of growth and $K_c$ is the half-saturation constant where $r(c)/r_{\max} = \frac{1}{2}$. We can obtain a single equation of logistic growth by 1) using the principle of mass conservation and 2) assuming that the half-saturation constant is sufficiently large relative to the initial concentration of supplied resources ($K_c \gg c(0)$). Under the principle of mass conservation, resources and yield-corrected abundances must sum to a constant total mass at any given time $B \equiv \frac{x(t)}{Y} + c(t)$. This constraint allows us to obtain a function for $c(t)$. The assumption $K_c \gg c(0)$ linearizes the growth rate $r(c) \approx r_{\max}c/K_c$. Using these two results, we obtain the following single differential equation

$$\frac{dx}{dt} = \frac{Br_{\max}}{K_c}x\left(1 - \frac{x}{YB}\right) \tag{S4a}$$

$$= \frac{x}{\tilde{\tau}}\left(1 - \frac{x}{\tilde{K}}\right) \tag{S4b}$$

where we have defined the timescale of growth and the carrying capacity, the terms of logistic growth, in mechanistic terms $\tilde{\tau} \equiv \frac{K_c}{Br_{\max}}$ and $\tilde{K} \equiv YB$. Therefore, a model of logistic growth is appropriate for microbial microcosm experiments provided that the half-saturation constant is sufficiently large relative to the supplied concentration of resources, a requirement that can, in principle, be manipulated by the experimenter.

### S4 Text: Deriving the stationary AFD for the SLM with a constant rate of migration

To determine the extent that continuous experimental manipulations of demography can alter the macroecological patterns captured by Eq. 1, we incorporate a migration term ($m_i$). By performing a change of variables for the relative abundance on the Langevin governing the dynamics of absolute abundance, we obtain

$$\frac{dx_i}{dt} = m_i + \frac{x_i}{\tau_i}\left(1 - \frac{x_i}{K_i}\right) + \sqrt{\frac{\sigma_i}{\tau_i}} x_i \xi_i(t) \tag{S5}$$

Where $m_i$ has dimensionless units and represents the number of individuals immigrating divided by the total number of individuals in the focal community, $m_i \equiv m_i^{(n)}/N$. It's clear that $\langle x_i \rangle \to K_i + m_i$ as $t \to \infty$. As in the main manuscript, we will use the Itô $\leftrightarrow$ Fokker-Planck correspondence to formulate a partial differential equation for the probability $P(x_i, t)$ that the $i$th species has abundance $x$ at time $t$ [5]

$$\frac{\partial P}{\partial t} = -\frac{\partial}{\partial x_i}\left[\left(m_i + \frac{x_i}{\tau_i}\left(1 - \frac{x_i}{K_i}\right)\right) P(x_i, t)\right] + \frac{\sigma_i}{2\tau_i}\frac{\partial^2}{\partial x_i^2}\left(x^2 P(x_i, t)\right) \tag{S6}$$

To solve for the stationary distribution $P^*(x_i) = \lim_{t\to\infty} P(x_i, t)$ we first set the left hand side of Eq.S6 to zero and rearrange

$$\left(m_i + \frac{x_i}{\tau_i}\left(1 - \frac{x_i}{K_i}\right)\right) P^*(x_i) = \frac{\sigma_i}{2\tau_i}\frac{\partial}{\partial x_i}\left(x_i^2 P^*(x_i)\right) \tag{S7}$$

Following the derivation in [6], we set $x_i^2 P^*(x_i) \equiv Q(x_i)$, obtaining

$$\left(\frac{m_i}{x^2} + \frac{1}{x_i \tau_i} - \frac{1}{K_i \tau_i}\right) Q(x_i) = \frac{\sigma_i}{2\tau_i} Q^{'}(x_i) \tag{S8}$$

which has the following solution

$$Q(x_i) = c \cdot \exp\left[-\frac{2}{\sigma_i x_i}\left(\tau_i m_i + \frac{x_i^2}{K_i}\right)\right] x_i^{2\sigma_i^{-1}} \tag{S9}$$

where $c$ is an arbitrary constant. Using our definition of $Q(x_i)$ and fixing $c$ by imposing $\int_0^1 dx_i P^*(x_i) = 1$, we solve the integral

$$\int_0^\infty P^*(x_i) dx_i = \int_0^\infty c \cdot \exp\left[-\frac{2}{\sigma_i x_i}\left(\tau_i m_i + \frac{x_i^2}{K_i}\right)\right] x_i^{2\sigma_i^{-1}-2} dx_i \tag{S10}$$

The solution to this integral can be solved using a known identity (Eq. 2.3.16.1 in [7]). We derived a solution for this known identity for the sake of completeness. First, to simplify the integral we define the following parameters: $p \equiv \frac{2x_i}{\sigma_i K_i}$, $q \equiv \frac{2\tau_i m_i}{\sigma_i}$, $\alpha \equiv 2\sigma_i^{-1} - 2$, and set $x_i \equiv Ce^{\theta}$. Using this last definition, we obtain $px_i + \frac{q}{x_i} = aCe^{\theta} + \frac{b}{C}e^{-\theta}$. We can then chose $pC = \frac{q}{C} \Rightarrow C = \sqrt{\frac{q}{p}}$, from which we obtain $px_i + \frac{q}{x_i} = p\sqrt{\frac{q}{p}}\left(e^{\theta} + e^{-\theta}\right) = 2\sqrt{pq}\cosh(\theta)$. Using this identity, we can solve the integral

$$\int_0^\infty \exp\left[-\left(px_i + \frac{q}{x_i}\right)\right] x_i^\alpha dx_i = \int_{-\infty}^\infty \exp\left[-2\sqrt{pq}\cosh\left(\theta\right)\right]\left(\frac{q}{p}e^\theta\right)^{\alpha+1} d\theta \tag{S11a}$$

$$= \left(\frac{q}{p}\right)^{\frac{1}{2}(\alpha+1)} \int_{-\infty}^\infty \exp\left[-2\sqrt{pq}\cosh\left(\theta\right)\right] e^{\theta(\alpha+1)} d\theta \tag{S11b}$$

$$= \left(\frac{q}{p}\right)^{\frac{1}{2}(\alpha+1)} \int_{-\infty}^\infty \exp\left[-2\sqrt{pq}\cosh(\theta)\right] \cdot \left[\cosh(\theta\left(\alpha+1\right)) + \sinh(\theta\left(\alpha+1\right))\right] d\theta \tag{S11c}$$

$$= 2\left(\frac{q}{p}\right)^{\frac{1}{2}(\alpha+1)} \int_0^\infty \exp\left[-2\sqrt{pq}\cosh(\theta)\right] \cdot \cosh(\theta\left(\alpha+1\right)) d\theta \tag{S11d}$$

$$= 2\left(\frac{q}{p}\right)^{\frac{1}{2}(\alpha+1)} B_{\alpha+1}(2\sqrt{pq}) \tag{S11e}$$

where $B$ is the modified Bessel function of the second kind. After replacing our variables, we obtain a solution for the constant of integration

$$c = \left[2\left(\tau_i m_i K_i\right)^{\frac{1}{2}\left(2\sigma_i^{-1}-1\right)} \cdot B_{\left(2\sigma_i^{-1}-1\right)}\left(\frac{4}{\sigma_i}\sqrt{\frac{\tau_i m_i}{K_i}}\right)\right]^{-1} \tag{S12}$$

which we use to arrive at a solution for the stationary PDF

$$P^*(x_i) = \left[2\left(\tilde{m}_i K_i\right)^{\frac{1}{2}\left(2\sigma_i^{-1}-1\right)} \cdot B_{\left(2\sigma_i^{-1}-1\right)}\left(\frac{4}{\sigma_i}\sqrt{\frac{\tilde{m}_i}{K_i}}\right)\right]^{-1} \cdot \exp\left[-\frac{2}{\sigma_i x_i}\left(\tilde{m}_i + \frac{x_i^2}{K_i}\right)\right] x_i^{2\sigma_i^{-1}-2} \tag{S13}$$

which is the predicted form of the AFD with migration. We can replace the product containing the migration term with a unitless compound parameter $\tilde{m}_i \equiv m_i \tau_i$. Using this distribution, we can then calculate the first and second moments of the PDF as

$$\begin{aligned}\langle x_i \rangle &= \int_0^\infty P^*(x_i)\, x_i\, dx_i \\ &= \left[2\left(\tau_i m_i K_i\right)^{\frac{1}{2}\left(2\sigma_i^{-1}-1\right)} \cdot B_{\left(2\sigma_i^{-1}-1\right)}\left(\frac{4}{\sigma_i}\sqrt{\frac{\tau_i m_i}{K_i}}\right)\right]^{-1} \\ &\quad \cdot \int_0^\infty \exp\left[-\frac{2}{\sigma_i x_i}\left(\tau_i m_i + \frac{x_i^2}{K_i}\right)\right] x_i^{2\sigma_i^{-1}-1} dx_i \\ &= \left(\tau_i m_i K_i\right)^{\frac{1}{2}} \frac{B_{\left(2\sigma_i^{-1}\right)}\left(\frac{4}{\sigma_i}\sqrt{\frac{\tau_i m_i}{K_i}}\right)}{B_{\left(2\sigma_i^{-1}-1\right)}\left(\frac{4}{\sigma_i}\sqrt{\frac{\tau_i m_i}{K_i}}\right)}\end{aligned}$$

$$
\begin{aligned}
\langle x_i^2 \rangle &= \int_0^\infty P^*(x_i)\, x_i^2 \, dx_i \\
&= \left[ 2\left(\tau_i m_i K_i\right)^{\frac{1}{2}\left(2\sigma_i^{-1}-1\right)} \cdot B_{\left(2\sigma_i^{-1}-1\right)}\left( \frac{4}{\sigma_i}\sqrt{\frac{\tau_i m_i}{K_i}} \right) \right]^{-1} \\
&\quad \cdot \int_0^\infty \exp\left[ -\frac{2}{\sigma_i x_i}\left( \tau_i m_i + \frac{x_i^2}{K_i} \right) \right] x_i^{2\sigma_i^{-1}} \, dx_i \\
&= \tau_i m_i K_i \frac{B_{\left(2\sigma_i^{-1}+1\right)}\left( \frac{4}{\sigma_i}\sqrt{\frac{\tau_i m_i}{K_i}} \right)}{B_{\left(2\sigma_i^{-1}-1\right)}\left( \frac{4}{\sigma_i}\sqrt{\frac{\tau_i m_i}{K_i}} \right)}
\end{aligned}
$$

From which we can derive the squared coefficient of variation

$$
\frac{\langle x_i^2 \rangle - \langle x_i \rangle^2}{\langle x_i \rangle^2} = \left( \frac{B_{\left(2\sigma_i^{-1}+1\right)}\left( \frac{4}{\sigma_i}\sqrt{\frac{\tau_i m_i}{K_i}} \right)}{B_{\left(2\sigma_i^{-1}\right)}\left( \frac{4}{\sigma_i}\sqrt{\frac{\tau_i m_i}{K_i}} \right)} \right)^2 - 1 \tag{S14}
$$

We ultimately want to test the feasibility of the sampling distribution, which we define as the convolution of the AFD and the probability of sampling $n^{\text{reads}}$ for a given species from a Poisson distribution out of $N^{\text{reads}}$ total reads.

$$
\begin{aligned}
P(n_i | K_i, \sigma_i, \tilde{m}_i, N) &= \int_0^\infty P^*(x_i | K_i, \sigma_i, \tilde{m}_i) P(n_i | N, x_i) \, dx_i \\
&= \left[ 2\left(\tilde{m}_i K_i\right)^{\frac{1}{2}\left(2\sigma_i^{-1}-1\right)} \cdot B_{\left(2\sigma_i^{-1}-1\right)}\left( \frac{4}{\sigma_i}\sqrt{\frac{\tilde{m}_i}{K_i}} \right) \right]^{-1} \\
&\quad \cdot \frac{N^{n_i}}{n_i!} \int_0^\infty \exp\left[ -\left( \frac{2\tilde{m}_i}{\sigma_i x_i} + \frac{2x_i}{\sigma_i K_i} + N x_i \right) \right] x^{2\sigma_i^{-1}-2+n_i} e^{-N x_i} \, dx_i \\
&= \frac{N^{n_i}}{n_i!} \left(\tilde{m}_i K_i\right)^{-\frac{1}{2}\left(2\sigma_i^{-1}-1\right)} \left( \frac{2\tilde{m}_i K_i}{2 + N\sigma_i K_i} \right)^{\frac{1}{2}\left(2\sigma^{-1}-1+n_i\right)} \\
&\quad \cdot \frac{B_{\left(2\sigma_i^{-1}-1+n_i\right)}\left( \sqrt{\frac{8\tilde{m}_i}{\sigma_i}\left( \frac{2}{\sigma_i K_i} + N \right)} \right)}{B_{\left(2\sigma_i^{-1}-1\right)}\left( \frac{4}{\sigma_i}\sqrt{\frac{\tilde{m}_i}{K_i}} \right)}
\end{aligned}
$$

While the terms describing the mean and squared CV are correct, they are unwieldy. To make progress, it is useful to reduce Eq. S13 at different parameter limits to examine its behavior as well as how it differs from the stationary PDF of the SLM in the absence of migration (Eq. S7). We start be identifying limiting forms of the Bessel function. In the limit $B_v(y) \sim \frac{1}{2}\Gamma(v)\left(\frac{y}{2}\right)^{-v}$ as $y \to 0$, which corresponds to $\tilde{m}_i \ll \left(\frac{4}{\sigma_i}\right)^2 K_i$, the moments of the distribution reduce to those obtained from the SLM without migration.

Contrastingly, as $y \to \infty$ the Bessel function can be approximated by the asymptotic expansion $B_v(y) \sim \sqrt{\frac{\pi}{2y}} e^{-y} \left[ 1 + \frac{4v^2-1}{8y} + \frac{\left(4v^2-1\right)\left(4v^2-9\right)}{2!(8y)^2} + \cdots \right]$, which corresponds to the high migration limit $\tilde{m}_i \gg \left(\frac{4}{\sigma_i}\right)^2 K_i$. Using the first term of this expansion, we arrive at the stationary PDF.

$$
P^*(x_i) = \sqrt{\frac{8}{\pi\sigma_i}} \left( \frac{\tau_i m_i}{K_i} \right)^{\frac{1}{4}} \cdot \exp\left[ \frac{2}{\sigma_i}\left( 2\sqrt{\frac{\pi m_i}{K_i}} - \frac{\tau_i m_i}{x_i} - \frac{x_i}{K_i} \right) \right] x_i^{2\sigma^{-1}-1} \tag{S15}
$$

This limiting form of the stationary PDF is, again, unwieldy, but it is clear from visual inspection that it does not resemble the gamma distribution. By taking the same limit of the moments of the AFD with migration, we obtain the following approximations

$$\begin{aligned}
\langle x_i \rangle &\approx \sqrt{\tilde{m}_i K_i}\,\frac{\frac{32}{\sigma}\sqrt{\frac{\tilde{m}_i}{K_i}} + 4(\frac{2}{\sigma})^2 - 1}{\frac{32}{\sigma}\sqrt{\frac{\tilde{m}_i}{K_i}} + 4(\frac{2}{\sigma} - 1)^2 - 1} \\
\frac{\left\langle x_i^2 \right\rangle - \left\langle x_i \right\rangle^2}{\left\langle x_i \right\rangle^2} &\approx \left( \frac{1 + \frac{4(2\sigma_i^{-1}+1)^2 - 1}{\frac{32}{\sigma_i}\sqrt{\frac{\tilde{m}_i}{K_i}}}}{1 + \frac{4(2\sigma_i^{-1})^2 - 1}{\frac{32}{\sigma_i}\sqrt{\frac{\tilde{m}_i}{K_i}}}} \right)^2 - 1
\end{aligned} \tag{S16}$$

### S5 Text: Obtaining the time-dependent AFD for the SLM

To contrast the effect of a constant rate of migration (i.e., chemostat) and migration as an initial condition (i.e., batch culture) we used the time-dependent probability distribution of abundances for the SLM with no migration ($P(x,t|x_0)$). To our knowledge, this solution was first derived by Schenzle and Brand and later rederived by Otunuga [8, 9]. Using the latter derivation, the time-dependent distribution is

$$\begin{aligned} P(x,t|x_0) = x^{2\sigma_i^{-1}-1} e^{-\frac{2x}{K_i\sigma_i}} \sum_{m=0}^{M} \Bigg[ \left(\frac{2}{K_i\sigma_i}\right)^{v_m} \frac{m! v_m}{\Gamma(v_m+m+1)} e^{-\lambda_n t} (x * x_0)^{-m} \\ \cdot L_m^{v_m}\left(\frac{2x}{K_i\sigma_i}\right) L_m^{v_m}\left(\frac{2x_0}{K_i\sigma_i}\right)\Bigg] \\ + A x^{2\sigma_i^{-1}-1} e^{-\frac{2x}{K_i\sigma_i}} \\ \cdot \int_0^\infty e^{-\lambda(\eta)t} h(\eta, x_0^{-1}) h(\eta, x^{-1}) d\eta \end{aligned} \tag{S17}$$

where $L_m^{v_m}$ is the Laguerre polynomial of degree $m$ and the summation occurs over the range $\sigma_i^{-1} - \frac{3}{2} \leq M \leq \sigma_i^{-1} - \frac{1}{2}$. The remaining terms of the function are

$$\lambda_m = \frac{m\sigma_i}{2\tau_i}\left(2\sigma_i^{-1} - 1 - \eta\right) \tag{S18a}$$

$$\lambda(\eta) = \frac{\sigma_i}{2\tau_i}\left(\left(\sigma_i^{-1} - \frac{1}{2}\right) + \eta^2\right) \tag{S18b}$$

$$h(\eta, y) = G_\eta y^{\sigma_i^{-1}} e^{\frac{1}{K_i\sigma_i y}} W_{\sigma_i^{-1}, i\eta}\left(\frac{1}{K_i\sigma_i y}\right) \tag{S18c}$$

$$G_\eta = \sqrt{\frac{\sigma_i K_i}{2\pi^2 A}\eta \sinh(2\pi\eta)\Gamma\left(\frac{1}{2} - \sigma_i^{-1} + i\eta\right)\Gamma\left(\frac{1}{2} - \sigma_i^{-1} - i\eta\right)} \tag{S18d}$$

$$v_m = 2\sigma_i^{-1} - 2m \tag{S18e}$$

$$A = \left[\left(\frac{\sigma_i K_i}{2}\right)^{2\sigma_i^{-1}-1} \Gamma\left(2\sigma_i^{-1} - 1\right)\right]^{-1} \tag{S18f}$$

where $W_{\kappa, i\eta}(\cdot)$ is the second solution to the Whittaker differential equation. To briefly summarize how this solution was obtained, the solution to the FPE was assumed to follow an eigenvalue form, from which a Kolmogorov Backward Equation was obtained. Using the stationary solution, a differential equation was obtained that reduced to the Kummer differential equation, from which a solution was obtained. Here we have used the solution of the Itô form of the SLM with migration, consistent with our derivation in S1.

### S6 Text: Simulating the SLM

Here we describe how sampling, both in terms of inoculation as well as sequencing, was implements in our SLM simulations as well as how the SLM simulations were performed. The process of sampling community members at the end of a transfer cycle can be modeled as multinomial sampling process, where we obtain a vector of initial abundances of all ASVs $\vec{n}^{(k)}(0)$

$$\Pr\left[\vec{n}^{(k)}(0)\right] = \begin{cases} \frac{N_{\mathrm{inoc}}!}{n_1!\cdots n_S!} x_{1,\mathrm{progenitor}}^{n_1} \cdots x_{S,\mathrm{progenitor}}^{n_S}, & \text{if } k = 0 \\ \frac{N^{(k)}(0)!}{n_1!\cdots n_S!} (x_1^{(k-1)}(T))^{n_1} \cdots (x_S^{(k-1)}(T))^{n_S}, & \text{if } k > 0 \end{cases} \tag{S19}$$

The vector of $S$ ASV abundances at the start of the $k$th transfer cycle ($\vec{n}^{(k)}(0)$) are drawn from the progenitor community for the first transfer cycle ($k = 0$) and are then drawn from the previous transfer at $T = 48$ h. for all subsequent transfer cycles ($k > 0$). The term $N_{\mathrm{inoc}}$ represents the number of individuals sampled from the progenitor community when each replicate community was initiated.

Similarly, the manipulation of abundances at the start of a transfer cycle due to migration can be modeled as a multinomial sampling process

$$\Pr\left[\vec{n}^{(k)}_{\mathrm{migration}}\right] = \begin{cases} \frac{N_{\mathrm{regional}}!}{n_1!\cdots n_S!} x_{1,\mathrm{progenitor}}^{n_1} \cdots x_{S,\mathrm{progenitor}}^{n_S}, & \text{Regional, } k \leq 12 \\ \frac{N_{\mathrm{global}}!}{n_1!\cdots n_S!} (x_{1,\mathrm{global}}^{(k-1)})^{n_1} \cdots (x_{S,\mathrm{global}}^{(k-1)})^{n_S}, & \text{Global, } k \leq 12 \end{cases} \tag{S20}$$

where $\vec{n}^{(k)}_{\mathrm{migration}}(0) = \vec{0}$ for $k > 12$ since at this point the migration manipulations have ceased.

Beyond migration, there are dependencies between the final abundance of an ASV and its abundance in the progenitor community that we would like to incorporate into our model. The set of carrying capacities $K_i$ was drawn from the parameters obtained by fitting Eq. 9 to the empirical MAD. The dependence between abundances in the progenitor community $\vec{x}_{\mathrm{progenitor}}$ and the empirical MAD were evaluated by performing logistic regression from `scikit-learn v0.22.1` [10]. Under the SLM the mean relative abundance is proportional to the carrying capacity, allowing us to write our logistic regression formula in terms of $K_i$

$$\Pr[K_i > 0 | x_{i,\mathrm{progenitor}}] \propto \left[1 + \exp(-\left(a + b * \log_{10} x_{i,\mathrm{progenitor}}\right))\right]^{-1} \tag{S21}$$

where $a$ is the intercept and $b$ is the slope of the regression.

From this relationship we can model the probability that an ASV has a non-zero carrying capacity $\Pr[K_i > 0 | x_{i,\mathrm{progenitor}}]$, allowing us to define the full distribution of carrying capacities as follows

$$K_i \sim \begin{cases} \text{Lognorm} & \text{with probability } \Pr[K_i > 0 | x_{i,\mathrm{progenitor}}] \\ 0 & \text{with probability } 1 - \Pr[K_i > 0 | x_{i,\mathrm{progenitor}}] \end{cases} \tag{S22}$$

The total number of ASVs with $K_i > 0$ was drawn from the distribution $S_{\mathrm{descendant}} \sim \text{Binomial}(S_{\mathrm{progenitor}}, 0.016)$, where $S_{\mathrm{progenitor}}$ is the number of ASVs in the progenitor community and 0.016 is the mean fraction of ASVs that were present in the assembled communities relative to the number of ASVs in the progenitor community.

Grafting these empirical considerations onto the time-dependent analytic solution of the SLM is not straightforward. Therefore, we elected to simulate the SLM. We used an approximation of the numerical solution of the SLM for the dynamics within each transfer cycle, incorporating above experimental details while requiring only two parameters: $\sigma$ and $\tau$, the values of the latter being constrained by the amount of growth that can possibly occur over 48 h. within a transfer cycle. To simulate the SLM, we derived a form of Eq. 1 so that the noise was additive instead of multiplicative by introducing the change of variable $q_i = \log(x_i)$, expanding $dq_i$ as a Taylor series, and using Itô's lemma.

$$\begin{aligned} dq_i &= \frac{\partial q_i}{\partial t}dt + \frac{\partial q_i}{\partial x_i}dx_i + \frac{1}{2}\frac{\partial^2 q_i}{\partial x_i^2}(dx_i)^2 && \text{(S23a)} \\ &= \frac{\partial q_i}{\partial t}dt + \frac{\partial q_i}{\partial x_i}\left(\frac{x_i}{\tau_i}\left(1-\frac{x_i}{K_i}\right)dt + \sqrt{\frac{\sigma_i}{\tau_i}}x_i dW(t)\right) \\ &\quad + \frac{1}{2}\frac{\partial^2 q_i}{\partial x_i^2}\left(\frac{x_i}{\tau_i}\left(1-\frac{x_i}{K_i}\right)dt + \sqrt{\frac{\sigma_i}{\tau_i}}x_i dW(t)\right)^2 && \text{(S23b)} \\ &= \left(\frac{\partial q_i}{\partial t} + \frac{\partial q_i}{\partial x_i}\frac{x_i}{\tau_i}\left(1-\frac{x_i}{K_i}\right) + \frac{\sigma_i}{2\tau_i}\frac{\partial^2 q_i}{\partial x_i^2}x_i^2\right)dt \\ &\quad + \sqrt{\frac{\sigma_i}{\tau_i}}\frac{\partial q_i}{\partial x_i}x_i dW(t) && \text{(S23c)} \end{aligned}$$

where $W(t)$ is a Wiener process. We then obtain the Langevin by dividing both sides by $dt$

$$\frac{dq_i}{dt} = \frac{1}{\tau_i}\left(1 - \frac{\sigma_i}{2} - \frac{e^{q_i}}{K_i}\right) + \sqrt{\frac{\sigma_i}{\tau_i}}\eta(t) \tag{S24}$$

We can approximate the numerical solution to Eq. S24 using the Euler–Maruyama method [5], where we simulate the discretized form of the equation, defining $\delta t$ as a single generation.

$$q_i(t+\delta t) = q_i(t) + \frac{1}{\tau_i}\left(1 - \frac{\sigma_i}{2} - \frac{e^{q_i(t)}}{K_i}\right)\delta t + \sqrt{\frac{\sigma_i \delta t}{\tau_i}}Z(t) \tag{S25}$$

where $Z(t) \sim \mathcal{N}(0,1)$. Within each transfer Eq. S25 was run seven times, corresponding to $\log_2(D^{-1}_{\mathrm{transfer}}) \approx 7$ generations. We set $\sigma$ as a constant due to the existence of Taylor's Law and $\tau$ as a constant due to the unknown nature of the distribution of growth rates. We used logarithmically spaced parameter values from the following ranges: $\sigma \in [0.01, 1.9]$ and $\tau \in [1.7, 6.9]$. The upper bound on the range of $\sigma$ was set by the observation that $\langle x_i \rangle = 0$ for $\sigma \leq 2$. The range on $\tau$ was set by the observation that the serial dilution factor sets the total number of generations per-transfer as $\log_2(D^{-1}_{\mathrm{transfer}}) \sim 7$, assuming exponential growth. This translates to a maximum generation time of $\tau_{\mathrm{max}} = 48\mathrm{h.}/7 \sim 6.9\mathrm{h}$. We identified a reasonable bound on the minimum generation time by noticing that prior research efforts have established that these communities have a maximum growth rate of $\sim 0.6\,\mathrm{h.}^{-1}$, translating to a minimum generation time of $\tau_{\mathrm{min}} \equiv 0.6^{-1} \approx 1.7\mathrm{h.}$ [1].

Simulated relative abundances were calculated from simulated true abundances, with the sampling process modeled as a multinomial process

$$\Pr\left[\vec{n}^{(k)}_{\mathrm{reads}}\right] = \frac{N_{\mathrm{reads}}!}{n_1!\cdots n_S!}(x_1^{(k)})^{n_1}\cdots(x_S^{(k)})^{n_S} \tag{S26}$$

The empirical total number of reads $N_{\mathrm{reads}}$ was used for all simulations (see Fig. S7 for visualization of the variation in sampling effort). Information about the statistical analysis can be found in the Materials and Methods.

### S7 Text: Inferring SLM parameters

Parameter combinations of $\tau, \sigma$ that produced simulated statistics which most closely matched statistics estimated from the experimental data were identified using Approximate Bayesian Computation. For analyses with a single statistic we selected the simulation iteration with the lowest Euclidean distance between the observed and simulated statistic. For analyses where a set containing multiple test statistics ($\vec{c}$) were considered, we used a weighted measure of Euclidean distance where the simulated values of each test statistic were scaled by their standard deviation to prevent the distance from being dominated by a single statistic with high variance [11].

$$d(\vec{c}_{\mathrm{obs}}, \vec{c}_{\mathrm{sim}}) = \left[ \sum_{g=1}^{G} \left( \frac{c_{\mathrm{sim}}^{(g)} - c_{\mathrm{obs}}^{(g)}}{\sqrt{\mathrm{Var}(c_{\mathrm{sim}}^{(g)})}} \right)^2 \right]^{\frac{1}{2}} \tag{S27}$$

This procedure was only used to identify parameters for the regional migration statistics and for the global migration $\Delta \ell$ statistics. Once the optimal values of $\tau, \sigma$ were identified they were used to perform $10^3$ simulations of our SLM model, generating a distribution of predicted summary statistics that was compared to the observed summary statistic.

Our rationale for using ABC was that it provided the flexibility necessary to investigate the properties of microbial communities, where the variation one observes can be modeled as an outcome of a disordered system (i.e., parameters such as carrying capacity being drawn from a specified distribution [12]) and 2) is in-part driven by the inherent stochasticity of sampling. This flexibility comes with the constraint that the effectiveness of ABC can be limited by the availability of informative statistics for a given model [13, 14]. In addition, the number of summary statistics presents its own issues, as it increases the dimensionality of the sampling space, resulting in an exponential decline in the probability of accepting a given simulation under ABC criteria [15]. However, it is unlikely that these issues shaped the results of this study. Whenever possible, we elected to examine patterns and quantities that are frequently used and tested in macroecological studies. When it was necessary to identify additional statistics to evaluate the macroecological outcomes of specific treatments, we scoured the literature for statistical tests that account for the standard error of an estimator (e.g., Fisher's Z-statistic for the difference in correlation coefficients, Eq. 13). Dimensionality is also unlikely to be a major contributor, as we used at most two summary statistics to infer our two parameters. However, given that the identification of treatment-specific statistics was vital for testing the effects of specific forms of migration, it is apparent that the appropriate summary statistic(s) for a given experiment are not always obvious *a priori* and that investigating a variety of summary statistics is a worthwhile effort if one chooses to proceed with ABC.

### Figures

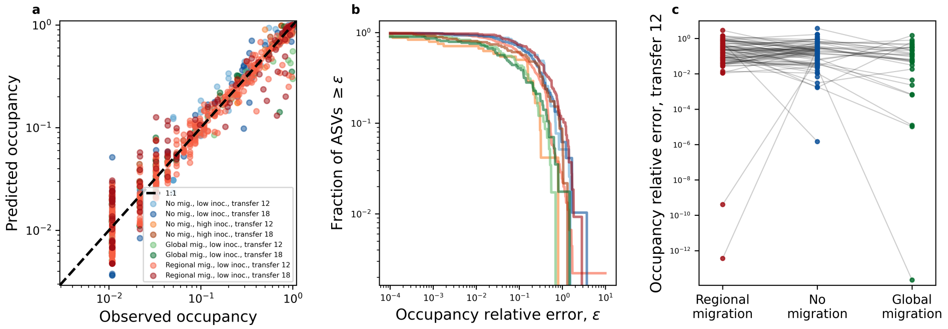


**Figure S1. The sampling form of the gamma distribution predicts ASV occupancy. a)** The fraction of replicates harboring a given ASV (i.e., occupancy) can be predicted using a form of the gamma distribution that accounts for sampling across experimental treatments and transfers. **b)** The distribution or relative errors exhibited a similar form across treatments and transfers, suggesting that the predictions of the SLM are broadly applicable. **c)** By comparing ASVs that were present in both migration and no migration treatments, we can see that the errors are generally similar between treatments, if only slightly higher in the no migration treatment.

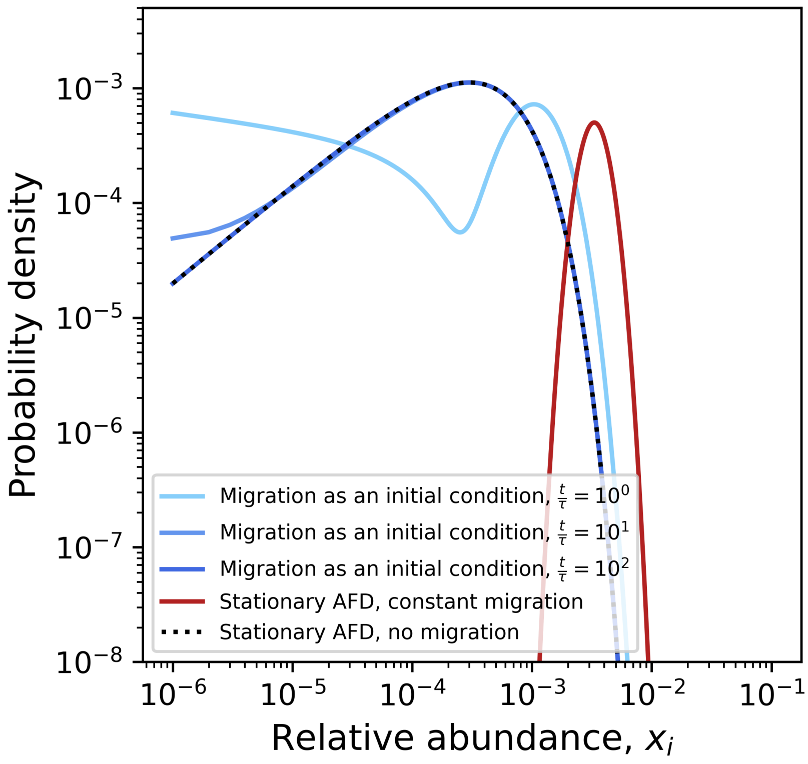


**Figure S2. The effect of migration on the AFD when modeled as a perturbation of initial conditions vs. a constant rate.** We examined how migration as a perturbation of initial conditions compared to a commonly assumed form of migration where it occurs at a constant rate per-unit time. The AFD of a form of the SLM with a constant rate of migration at stationarity was derived (Eq. S13 in S4 Text) and the time-dependent solution of the SLM (Eq. 13) was obtained from a prior study (S5 Text). The following parameters were used: $K_i = 10^{-3}$, $\sigma_i = 0.7$, $\tau = 1$, and $x_i(0) = m_i = 10^{-2}$. We have rescaled time using the timescale of growth to arrive at a dimensionless parameter $\frac{t}{\tau}$. The AFD with no experimentally-imposed migration is represented by Eq. 2 (i.e., a gamma distribution).

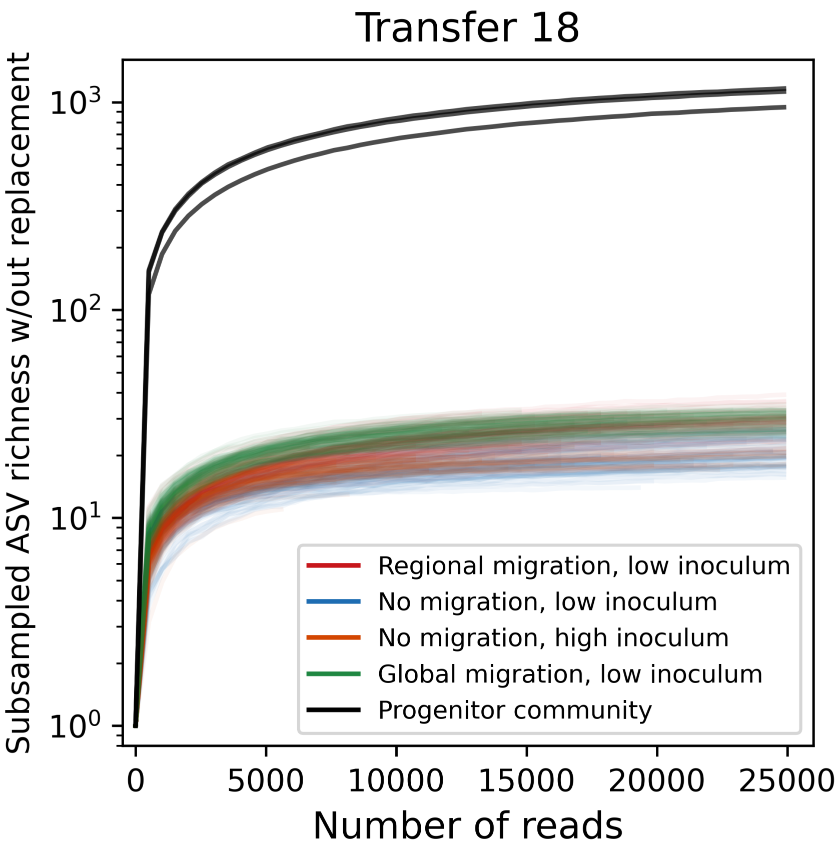


**Figure S3. Rarefaction curves for each treatment.** Rarefaction curves demonstrate how the richness (# ASVs) is 100-fold lower in descendant communities relative to the progenitor.

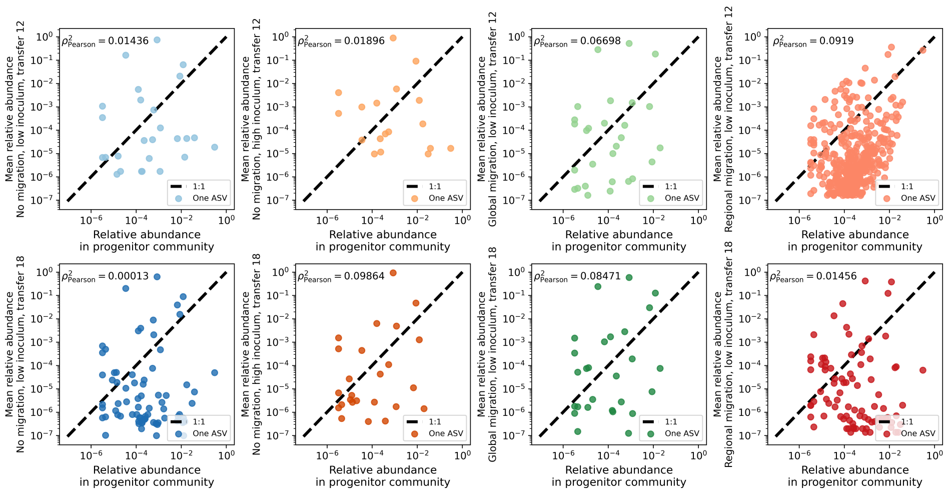


**Figure S4. Progenitor abundance vs. assembled mean abundance.** There is effectively zero correlation between the relative abundance of an ASV in the progenitor community and its mean abundance among descendant communities for all treatments at both transfers 12 and 18.

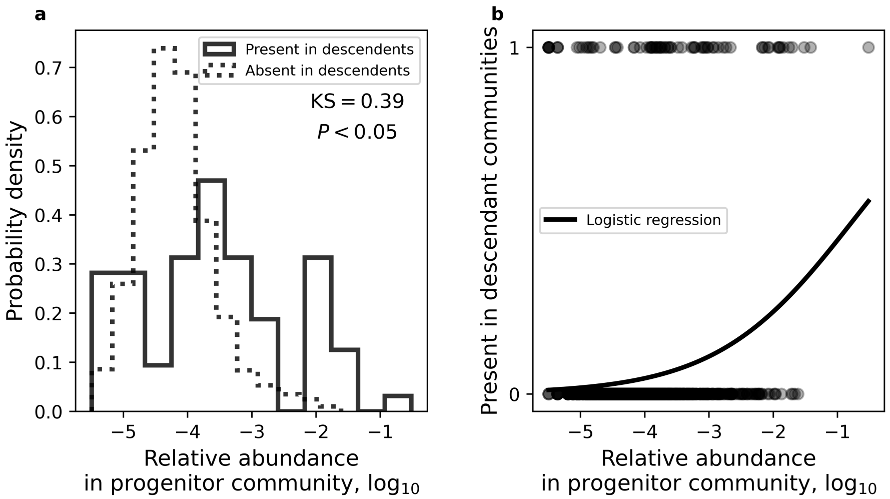


**Figure S5. Presence in descendent as function of progenitor abundance. a)** An ASV is more likely to be present in the descendant communities if it has a higher relative abundance in the progenitor. **b)** This result implies that probability that an ASV has a non-zero carrying capacity is a function of its progenitor abundance, a relationship that can be modeled as a logistic regression.

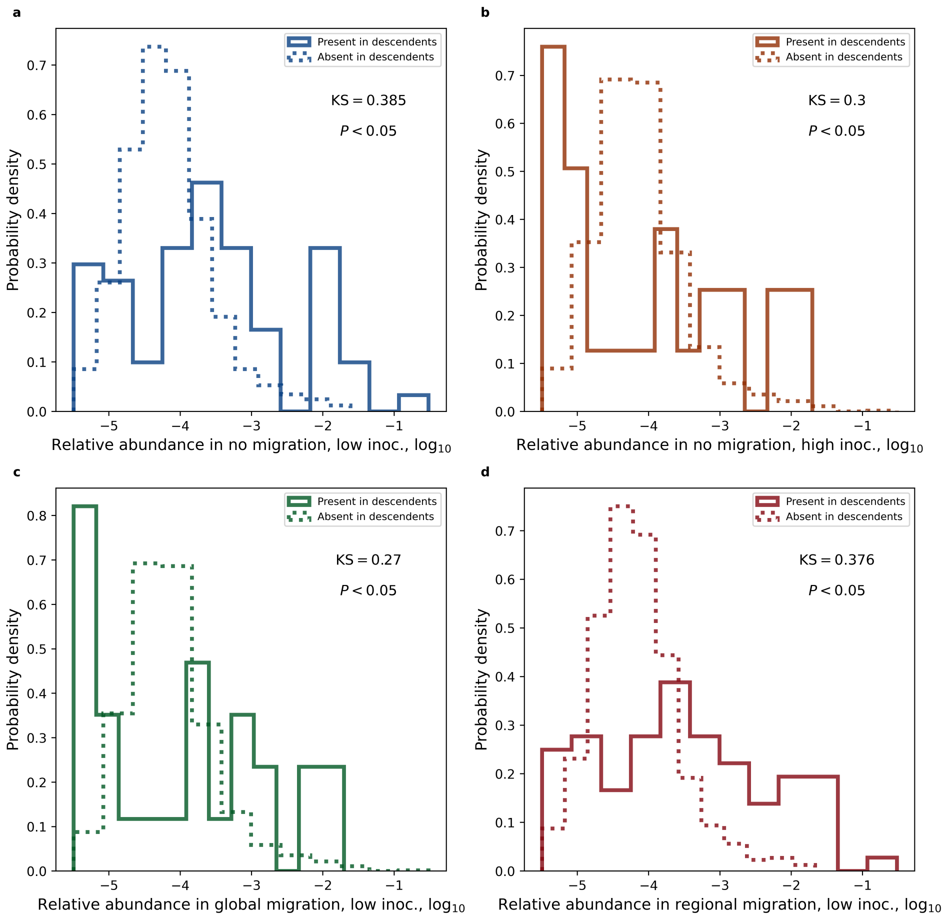


**Figure S6. The distribution of abundances in the progenitor community is consistent across treatment status. a-d)** The abundances of ASVs in the progenitor that are present in a given treatment are consistently shifted to the right, implying that the probability of an ASV surviving is conditional on its initial abundance. Permutation-based two-sample Kolmogorov–Smirnov tests were performed for each treatment.

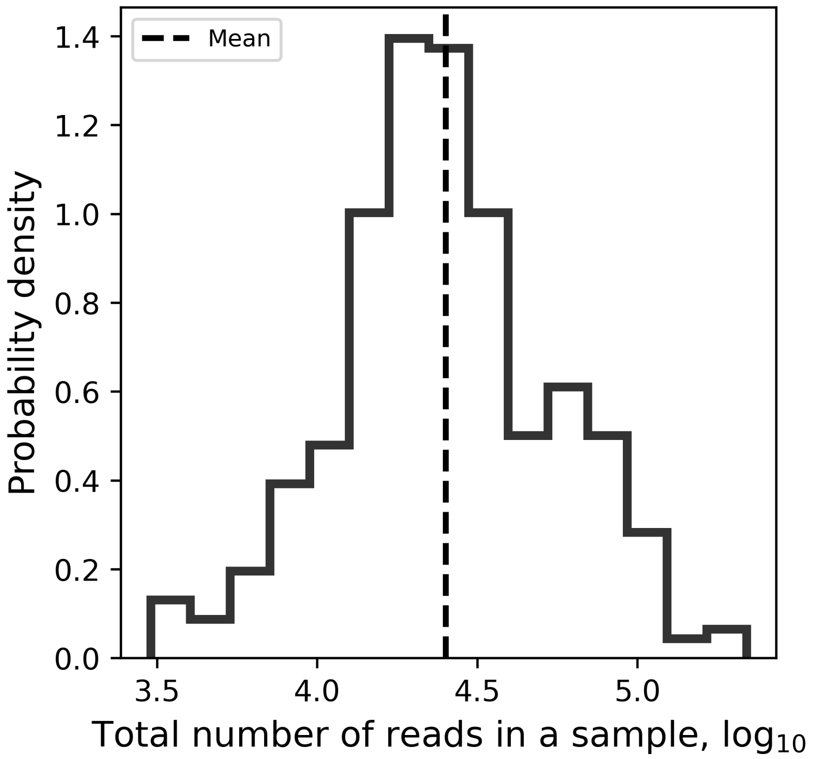


**Figure S7. The distribution of total read counts.** In our simulations the generation of reads from relative abundances was done as a multinomial sampling process, where the total number of reads of a given replicate at a given time point was drawn from the empirical distribution of total read counts.

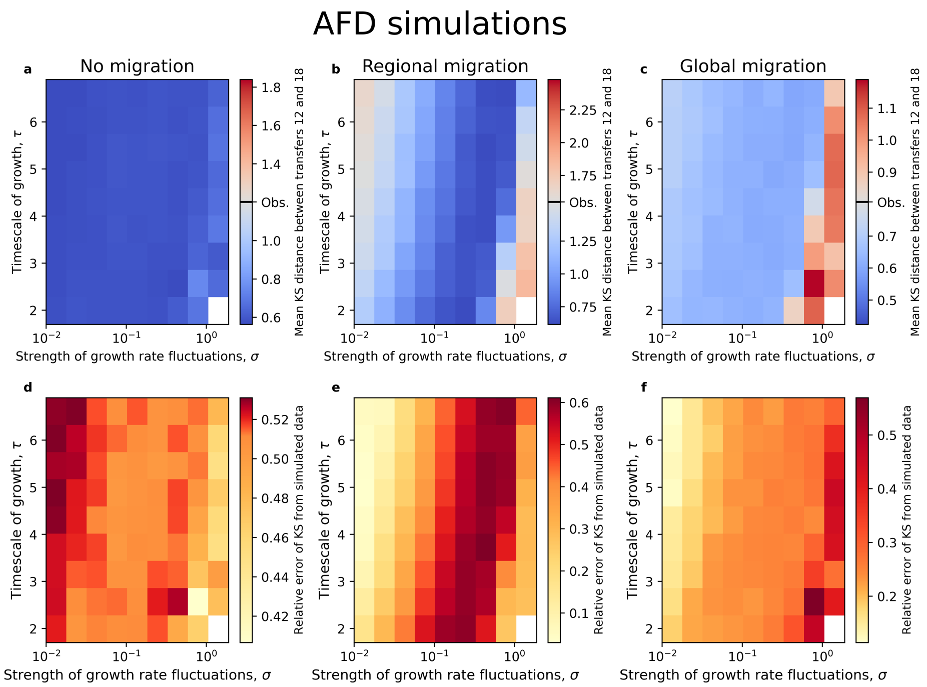


**Figure S8. AFD simulations**. **a-c)** The KS distance (KS) between AFDs from transfers 12 and 18 for all migration treatments across a parameter grid. **d-f)** Using the empirical KS distance, we obtained the mean relative error for each simulation. We see that the error of our predictions systematically tends to cluster for certain parameter regimes, particularly for regional migration. This pattern suggests that there are adjacent parameter regimes where the SLM adequately performs. For each parameter combination 100 iterations were performed.

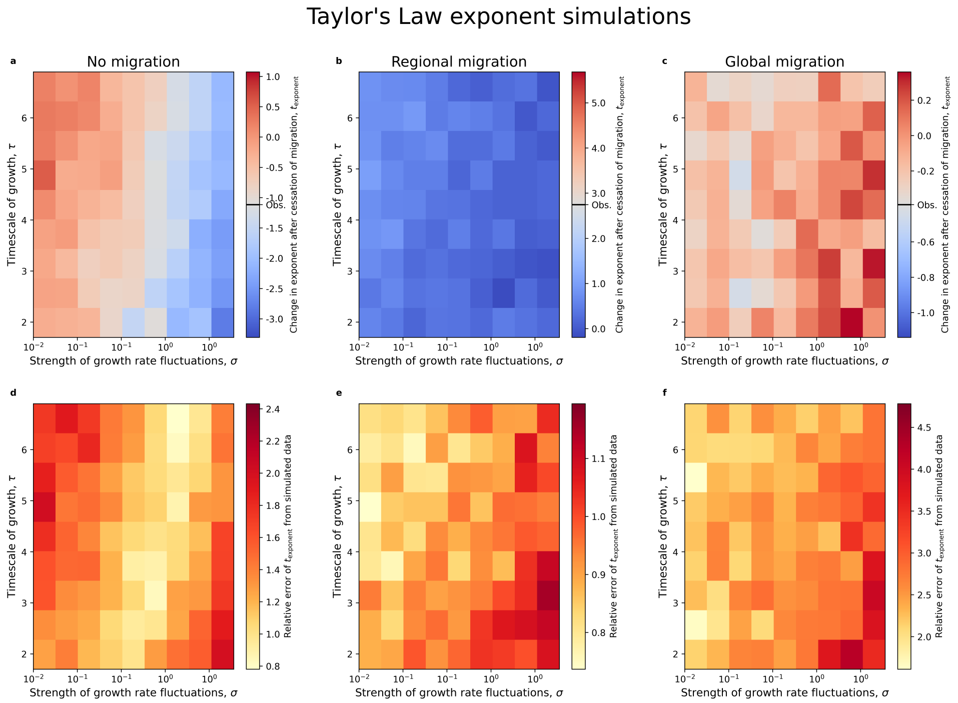


**Figure S9. Taylor's Law exponent simulations.** The equivalent analysis as shown in Fig. S8 for the exponent of Taylor's Law.

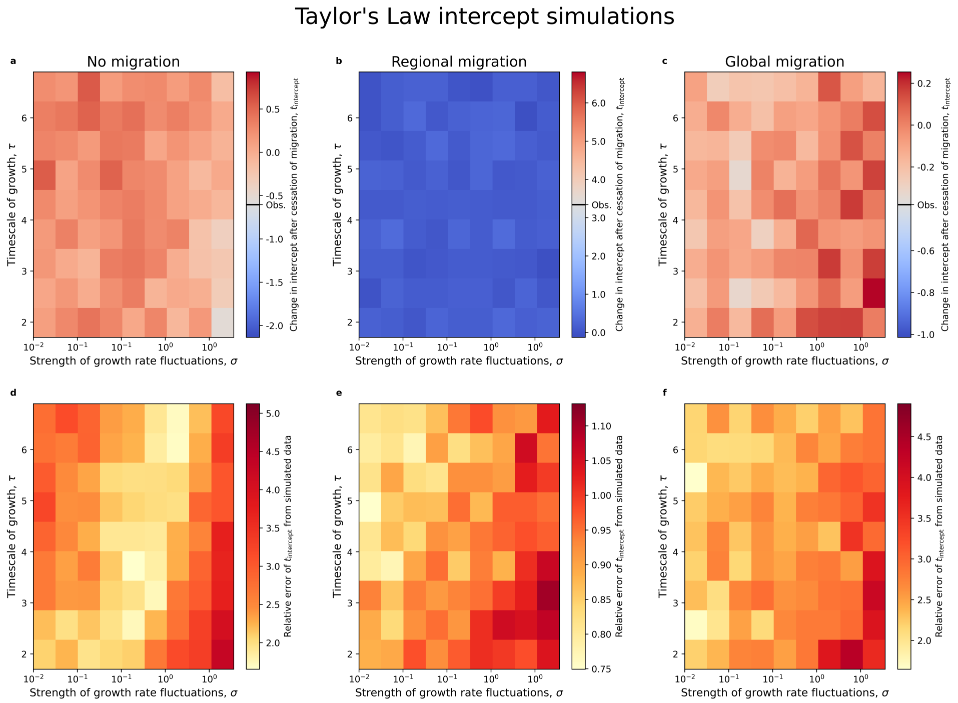


**Figure S10. Taylor's Law intercept simulations.** The equivalent analysis as shown in Fig. S8 for the intercept of Taylor's Law. Similar to Fig. S9 the errors tend to cluster together for regional migration, implying that the SLM is performing adequately for adjacent combinations of parameter regimes.

Regional migration statistics

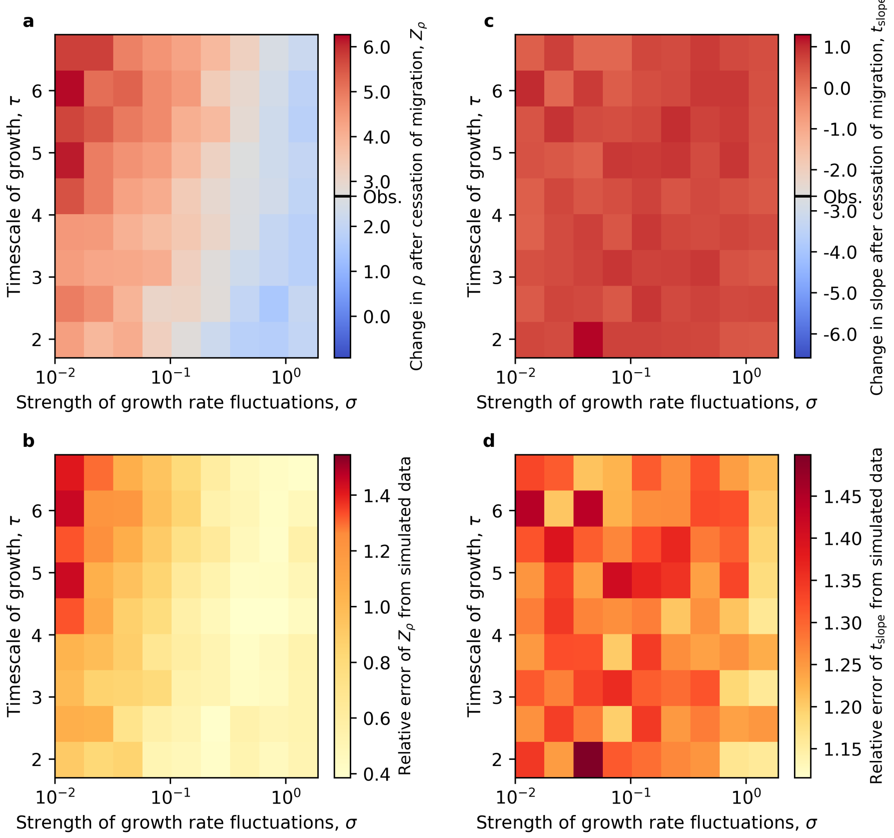


**Figure S11. Simulations of regional migration statistics.** The equivalent analysis as shown in Fig. S8 for statistics that capture the directional change in abundance caused by regional migration. The parameter regimes with lowest error correspond to the regions with lowest error for our regional migration Taylor's Law analyses (Fig. S9, S10).

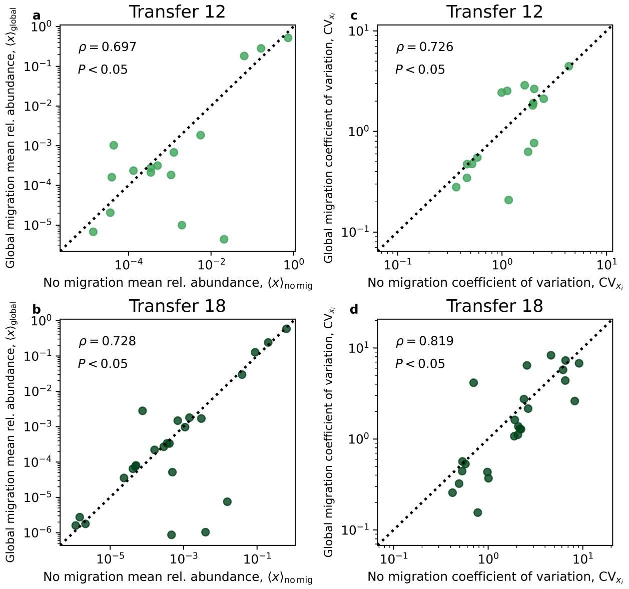


**Figure S12. Global migration does not alter the MAD nor the distribution of CVs.** Similarly to the regional migration patterns examined, we evaluated whether a change in correlation occurred for the global migration treatment for the paired MAD and distribution of CVs. Within each time point for each measure, the strength of the correlation of significantly greater than zero. However, the change in correlation $Z_{\rho}$ was not significant for both cases, a result that is consistent with simulation results.

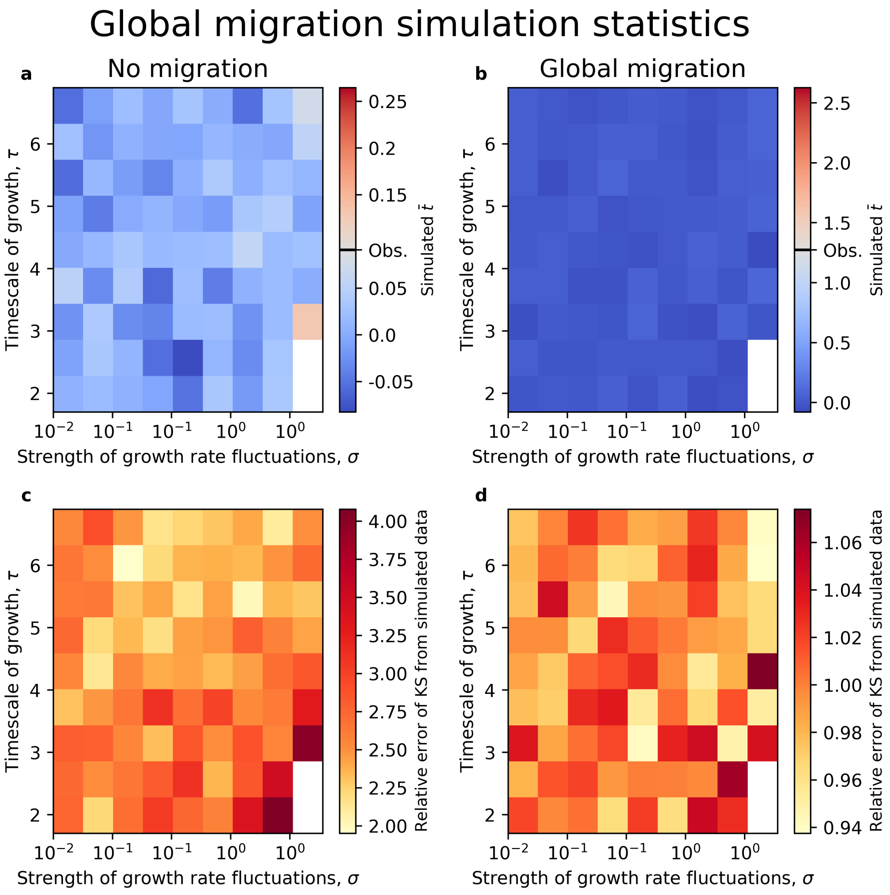


**Figure S13. Simulations of global migration statistics.** The equivalent analysis as shown in Fig. S8 for statistics that capture the change in the fluctuations around the typical abundance caused by global migration.

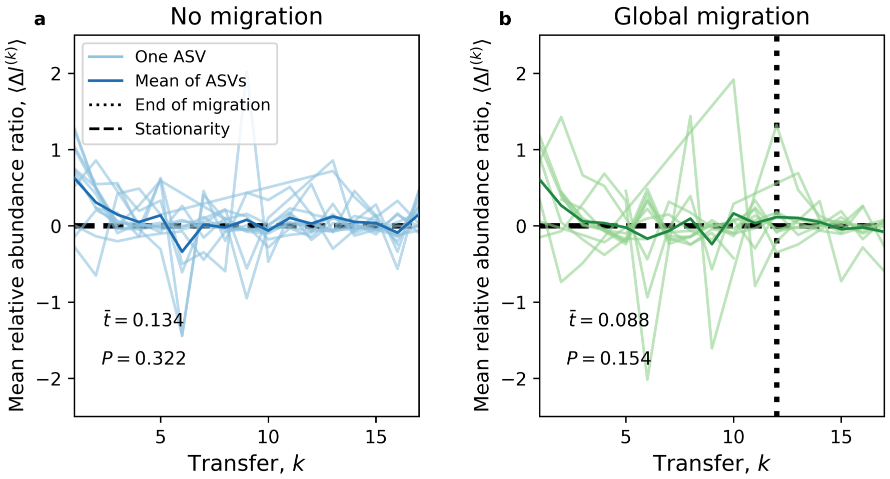


**Figure S14. Mean change in $\Delta\ell$ under global and no migration.** For both **a)** no and **b)** global migration the mean of $\Delta\ell$ is initially higher than the stationary value of zero, though the mean relaxes to zero by transfer six for both treatments and does not appear to change after the cessation of global migration. This result is consistent with predicted consequences of global migration.

### Tables

| Migration treatment | Inoculation | Transfer(s) | # sequenced communities |
|---|---|---|---|
| No migration | Low | 12 | 20 |
| | | 18 | 92 |
| | | 1-11, 13-17 | 20 |
| No migration | High | 12 | 4 |
| | | 18 | 93 |
| Regional | Low | 12 | 92 |
| | | 18 | 92 |
| | | 1-11, 13-17 | 8 |
| Global | Low | 12 | 93 |
| | | 18 | 93 |
| | | 1-11, 13-17 | 3 |

**Table S1.** The number of replicate communities sequenced for a given treatment at a given transfer.

| Transfer regime | Inoculation | % communities in a given attractor | |
|---|---|---|---|
| | | Alcaligenaceae | Pseudomonadaceae |
| No migration | Low | 70.7 | 29.3 |
| No migration | High | 0.0 | 100.0 |
| Regional | Low | 4.4 | 95.6 |
| Global | Low | 100.0 | 0.0 |

**Table S2.** The percent of communities belonging to a given attractor for each migration treatment, as described in [1].
